# aCPSF1 in synergy with terminator U-tract dictates archaeal transcription termination efficiency via the KH domains recognizing U-tract

**DOI:** 10.1101/2021.05.26.445813

**Authors:** Jie Li, Lei Yue, Wenting Zhang, Zhihua Li, Bing Zhang, Fangqing Zhao, Xiuzhu Dong

## Abstract

Recently, aCPSF1 was reported to function as the long-sought global transcription termination factor of archaea, while the working mechanism remains elusive. This work, through analyzing transcript-3′end-sequencing data of *Methanococcus maripaludis*, found positive correlations of both the terminator uridine(U)-tract and aCPSF1 with hierarchical transcription termination efficiencies (TTEs) at the genome-wide level. *In vitro* assays determined that aCPSF1 specifically binds to the terminator U-tract with U-tract number-related binding abilities, and *in vivo* assays demonstrated the two are indispensable in dictating high TTEs, revealing that aCPSF1 and the terminator U-tract in synergy determine high TTEs. The N-terminal KH domains equip aCPSF1 of specific binding to terminator U-tract and the *in vivo* aCPSF1-terminator U-tract synergism; aCPSF1’s nuclease activity was also required for TTEs. aCPSF1 also functioned as back-up termination for transcripts with weak intrinsic terminator signals. aCPSF1 orthologs from Lokiarchaeota and Thaumarchaeota exhibited similar U-tract synergy in dictating TTEs. Therefore, aCPSF1 and the intrinsic U-rich terminator could work in a noteworthy two-in-one termination mode in Archaea, which could be widely employed by archaeal phyla; using one factor recognizing U-rich terminator signal and cleaving transcript 3′-end, the archaeal aCPSF1-dependent transcription termination could display a simplified archetypal mode of the eukaryotic RNA polymerase II termination machinery.

## Introduction

Transcription termination is an essential and highly regulated process in all forms of life, which not only determines the accurate 3′-end boundary of a transcript and transcription related regulatory events, but also is important in shaping programmed transcriptomes of the living organisms (Peters et al., 2011; Porrua et al., 2016; Porrua and Libri, 2015; Ray-Soni et al., 2016; Yue et al., 2020). Highly controlled transcription termination, which prevents read-through resulted undesired increases in downstream coding regions and accumulation of antisense transcripts, can be particularly important in prokaryotes because of the gene densely packed genomes (Peters et al., 2012; Yue et al., 2020).

Researches have indicated that bacteria primarily employ two transcription termination mechanisms, Rho-dependent and -independent (intrinsic). In the former, the RNA translocase Rho, via recognizing a cytosine-rich sequence in nascent transcripts, dissociates the processive transcription elongation complex (TEC) based on its ATPase activity; while the intrinsic termination merely depends on a nascent RNA structure with a 7–8 base-paired hairpin followed by a run of uridines (Us) (Gusarov and Nudler, 1999; Peters et al., 2011; Porrua et al., 2016; Ray-Soni et al., 2016). A bacteria-like intrinsic termination mechanism that depends a U-stretch is also found in the eukaryotic RNA polymerase (RNAP) III (Nielsen et al., 2013; Orioli et al., 2011). Distinctively, transcription termination of the eukaryotic RNAP II, which transcribes mRNAs and non-coding RNAs, usually involves a transcript 3′-end processing event, in which the cleavage and polyadenylation complex (CPF/CPSF), under assistance of the accessory cleavage factors CFIA and CFIB, recognizes the termination signal, a poly(A) site at transcript 3′-end, cleaves at downstream of the nascent RNA, followed by polyadenylation for mRNA maturation, and triggers RNAP II dissociation for transcription termination (Baejen et al., 2017; Eaton et al., 2018; Grzechnik et al., 2015; Kim et al., 2004; Kuehner et al., 2011; Larochelle et al., 2018; Porrua et al., 2016).

Compared to bacteria and eukaryotes, knowledge of the transcription termination mechanisms and the related regulatory processes in the third form life, Archaea, is much lagging behind (Dar et al., 2016a; Maier and Marchfelder, 2019). Archaea represent a primary domain of cellular life and are phylogenetically more closely related to eukaryotes than bacteria (Eme et al., 2017; Williams et al., 2020; Zaremba-Niedzwiedzka et al., 2017). Specifically, they employ a eukaryotic RNAP II homolog, archaeal RNAP (aRNAP) (Werner and Grohmann, 2011), but have compact genomes with short intergenic regions (IGRs) and usually co-transcribed polycistronic operons, highlighting the importance of a controllable transcription termination. Earlier studies suggested that resembling to the bacterial intrinsic termination, transcription termination of aRNAP could depend merely on a short U-rich sequence at the transcript 3′-end without strictly requiring an upstream hairpin structure (Hirtreiter et al., 2010; Maier and Marchfelder, 2019; Santangelo et al., 2009; Santangelo and Reeve, 2006; Spitalny and Thomm, 2008; Thomm et al., 1994). Recently, through genome-level transcript 3′-end sequencing (Term-seq), the U-rich sequences preceding transcription termination sites (TTSs) without preceding hairpin structures were identified to be overrepresented in transcripts of four representative archaeal species, namely, *Methanosarcina mazei*, *Sulfolobus acidocaldarius*, *Haloferax volcanii* and *Methanococcus maripaludis* (Berkemer et al., 2020; Dar et al., 2016b; Yue et al., 2020). Therefore, the overrepresented U-tract sequences at the transcript 3′-ends are assumed as the intrinsic termination signals of archaea and not strictly requiring an upstream hairpin structure sets archaeal termination apart from the mechanism employed in bacteria (Maier and Marchfelder, 2019).

The protein factors that mediate archaeal transcription termination have just been reported in recent years. The *Thermococcus kodakarensis* Eta (<u>e</u>uryarchaeal termination <u>a</u>ctivity) has been reported to transiently engage the TEC and lead to stalled TEC released from damaged DNA lesions, thus resembling the bacterial Mfd termination factor and functioning specifically in response to DNA damage (Walker et al., 2017). Most recently, aCPSF1, also named FttA (Factor that Terminates Transcription in Archaea), is determined as a transcription termination factor of Archaea because it could competitively disrupt the processive TEC at normal transcription elongation rate and implement a kinetically competitive termination dependent on both the stalk domain of RNAP and the transcription elongation factor Spt4/5 *in vitro* (Sanders et al., 2020). aCPSF1 is affiliated within the β-CASP ribonuclease family, and is ubiquitously distributed in all archaeal phyla (Phung et al., 2013; Yue et al., 2020). Initially, it was assumed to function in RNA maturation and turnover of Archaea (Clouet-d’Orval et al., 2015), and endoribonuclease activities were identified for three aCPSF1 orthologs *in vitro* (Levy et al., 2011; Phung et al., 2013; Silva et al., 2011), with one also exhibiting 5′–3′ exoribonuclease activity (Phung et al., 2013). Our recent study reported that aCPSF1, depending on its ribonuclease activity, controls *in vivo* transcription termination at the genome-wide level and ensures programmed transcriptome in *M. maripaludis*, and its orthologs from the distant relatives, *Lokiarchaeota* and *Thaumarchaeota*, perform the same function in termination (Yue et al., 2020). However, although an *in vitro* enzymatic assay showed that aCPSF1 primarily cleaves endoribonucleolytically downstream of a U-rich motif that precedes the identified TTSs (Yue et al., 2020), some open questions remain, such as (i) whether the aCPSF1-dependent and the U-tract terminator-based intrinsic terminations are two independent mechanisms, or the two in fact work synergistically in archaea, or (ii) if the aCPSF1-dependent mode just serves as a backup mechanism for the genes/operons containing less-efficient intrinsic termination signals (Sanders et al., 2020; Wenck and Santangelo, 2020); (iii) what the exact sequence motifs are that aCPSF1 recognizes, and (iv) whether, like the eukaryotic multiple subunits composed termination complex, aCPSF1 also needs others in recognizing the termination signals.

In the present work, via an intensive analysis of the Term-seq data in *M. maripaludis,* we first systematically evaluated the correlations of the transcription termination efficiency (TTE) for each identified TTS with both the cis-element U-tract terminator and the trans-action termination factor aCPSF1. Further, in combination with molecular and genetic validations, we determined that aCPSF1 and the terminator U-tract synergistically dictate high TTEs. The *in vitro* and *in vivo* assays together demonstrated that the aCPSF1 N-terminal K homolog (KH) domains specifically recognize and bind to the terminator U-tract. Therefore, the archaeal termination factor aCPSF1 could accomplish the U-tract terminator recognition and transcript 3′-end cleavage by itself, and the factor-dependent transcription termination can be the primary mechanism used by Archaea.

## Results

### Term-seq reveals a positive correlation between the TTSs preceding four-uridine (U4) tract numbers and the transcription termination efficiency in *M. maripaludis*

Attempting to evaluate the specific termination signals recognized by the newly identified transcription termination factor aCPSF1 and its role in dictating the *in vivo* transcription termination efficiencies (TTEs), we intensively reanalyzed the genome-wide Term-seq data of *M. maripaludis* (Yue et al., 2020). By following the stringent filtration workflow of Transcription Termination Site (TTS) definition and to preclude identifying sites derived from stale RNA processing or degradation products, TTSs were all assigned within 200 nt downstream the stop codon of a gene to maximally enrich the authentic TTSs near the gene 3′-ends and only sites that appeared in both of the two biological replicates with significant coverage (see Methods) were analyzed. In total, 2357 TTSs, including the previously identified 998 primary and 1359 newly identified secondary TTSs, were obtained (Datasheet S1). Multiple consecutive TTSs were found in more than 50% of transcription units (TUs) of *M. maripaludis* (Figure 1- figure supplement 1), which could produce multi-isoforms of a transcript with varying 3′-UTRs, as that found in *M. mazei* and *S. acidocaldarius* (Dar et al., 2016a). Nevertheless, compared with the primary TTSs, which have the highest reads among the Term-seq captured transcript 3′-ends of each TU, much lower median reads abundances, TTEs and motif scores were identified for the secondary TTSs (Figure 1- figure supplement 2). This indicates that TUs are mainly terminated at the primary TTSs, which were therefore used for further investigation.

**Figure 1.**
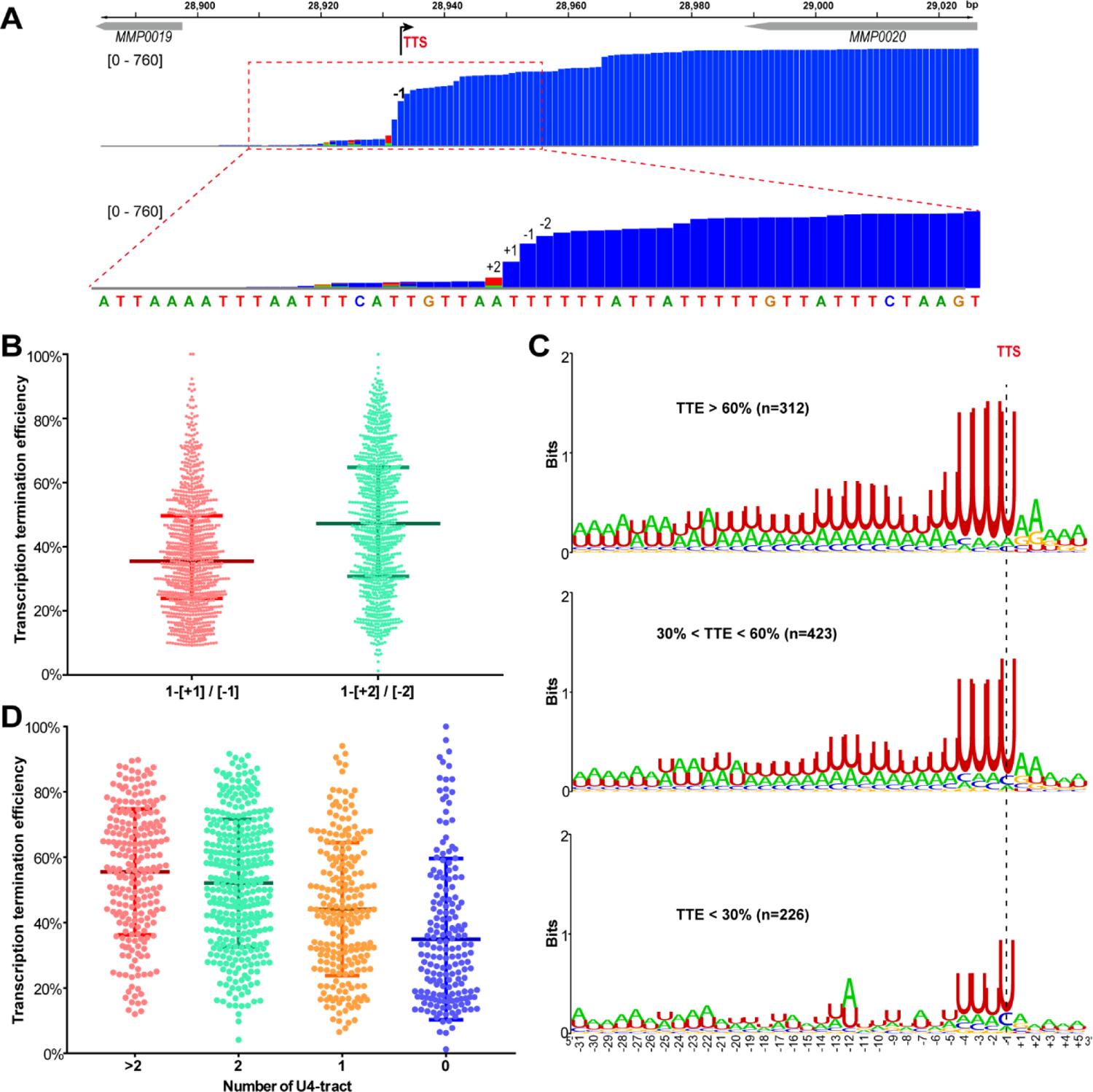
A positive correlation is observed between the terminator U4-tract numbers and transcription termination efficiencies (TTEs) among the TUs of *M. maripaludis*. (*A*) A representative Term-seq mapping map of *MMP0020* showing a stair-like descending pattern of the sequencing reads at four nucleotides that flank the identified transcription termination site (TTS, −1 site indicated by bent arrow). The magnified mapping region (red dotted frame) shows reads decreasing from −2 nt (upstream) to +2 (downstream) of the TTS in a step-wise manner. The chromosome locations of genes are indicated at the top, and the Term-seq reads mapping heights are shown in brackets. *(B)* Box-plot diagrams showing the TTE statistics of 998 transcripts that were calculated based on the reads ratio of nts +1 to −1 ([+1]/[−1]), and that of the nts +2 to −2 reads ([+2]/[−2]) respectively up- and down-stream the primary TTSs. Between the upper and lower lines are TTEs of 50% of transcripts, and the line in the middle represents the TTE median. *(C)* Logo representations of the terminator motif signatures in three groups of transcripts having different TTEs (>60%, >30% and <60%, <30%). The transcript numbers of each group are indicated inside parentheses. *(D)* Box-plot diagrams showing the statistics of TTE values among the four groups of terminators that carry various numbers of U-tracts. The diagram representations are the same as in (*B*). The significance statistical analysis for the TTEs of the four groups determined by Wilcoxon rank sum test were shown in Table S3.

**Figure 1- figure supplement 1.**
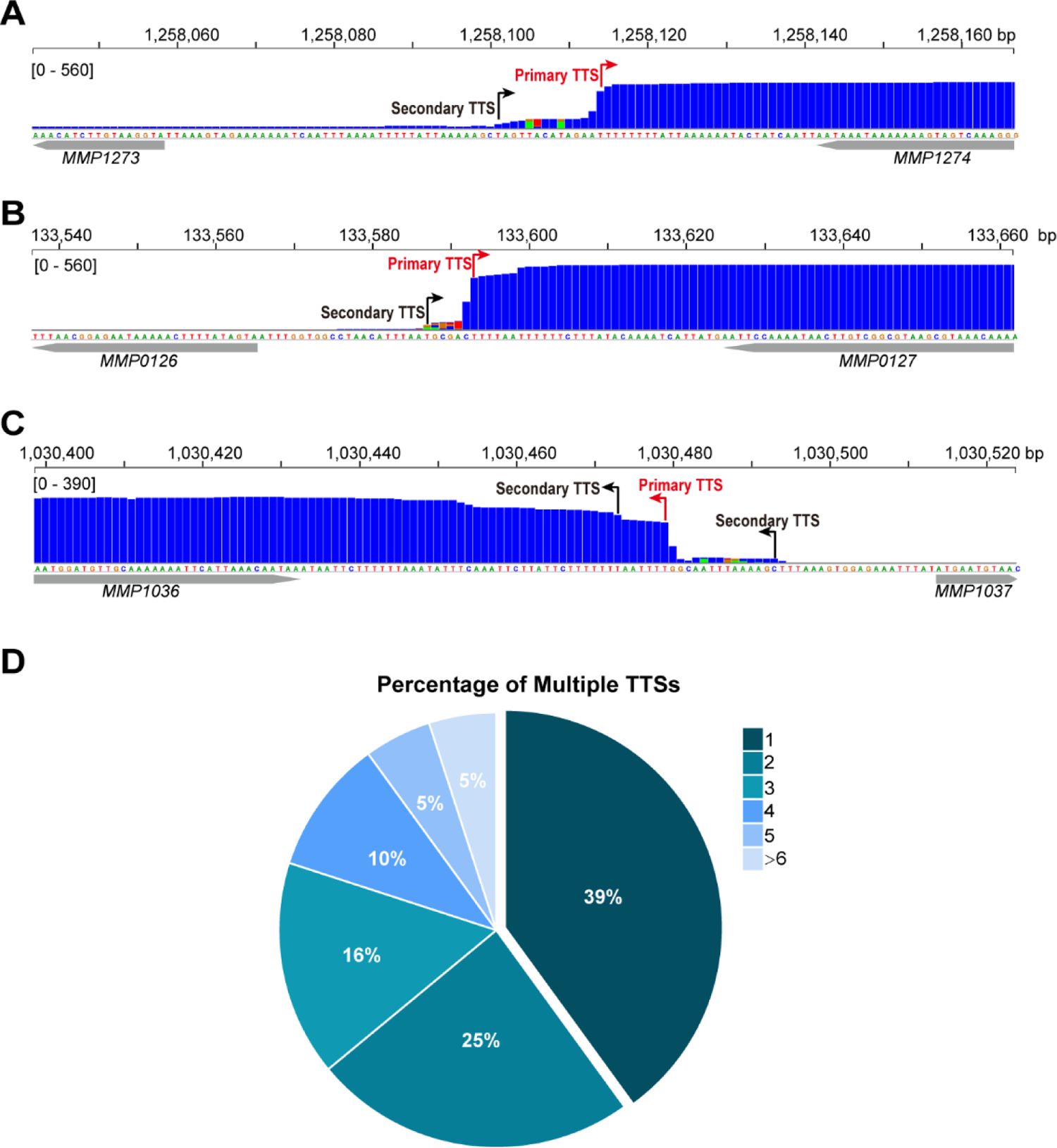
Multiple TTSs in one transcript identified by Term-seq in *M. maripaludis*. (A, B, C) Representative Term-seq mapping maps showed multiple TTSs in the *MMP1274* (A), *MMP0127* (B) and *MMP1036* (C) transcripts. The primary and secondary TTSs are indicated by red and black arrows, respectively. (D) Pie chart shows the percentages of TUs having 1 to >6 multiple TTSs identified.

Sequence analysis on the 961 primary TTSs of encoding TUs found a featured terminator motif, a 23 nt U-tract with the four consecutive uridine nucleotides (U4) that are most proximal to the TTS having the highest matching. To evaluate the contribution of the U-tract sequence preceding TTSs to transcription termination in *M. maripaludis*, we first defined and calculated transcription termination efficiency (TTE) of each TU. After inspecting the whole-genome Term-seq mapping file, a step-wise decreasing pattern in the mapping reads was observed at four nucleotides (nts) between sites +2 and −2 relative to the TTS (−1 nt) in vast majority of the primary TTSs (Figure 1A and Figure 1- figure supplement 3), i.e., transcription is terminated in a step-wise manner at the four nucleotides, which was therefore defined as the TTS quadruplet. Through pair-wisely comparing the reads of each nt in a TTS quadruplet, the maximal decrease was found between sites −2 nt (upstream) and +2 nt (downstream) relative to TTS in the majority of TTSs (Figure 1B). Thus, the reads ratio between −2 and +2 nts was used as the measurement of TTE, which was calculated based on “TTE=1− [+2] / [−2]”, where [−2] and [+2] represent the read abundances at −2 nt and +2 nt in Term-seq data, respectively.

**Figure 1- figure supplement 2.**
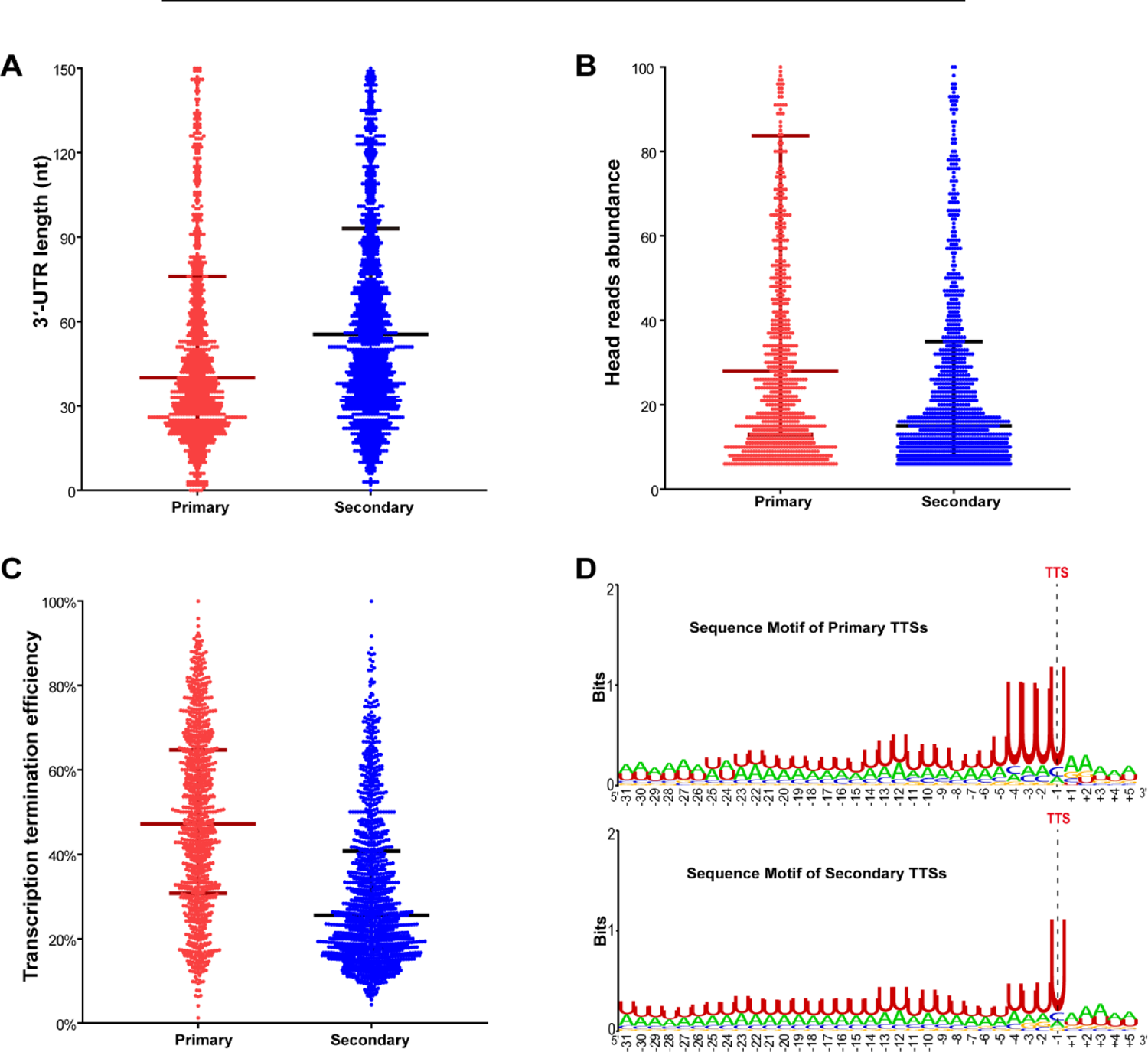
Features of primary and secondary TTSs determined in *M. maripaludis*. (A, B, C) Box-plot diagrams show the statistics of the 3′-UTR length (A), head read abundance (B), and transcription termination efficiencies (TTEs) (C) of the 998 primary and 1359 secondary TTSs, respectively. (D) Logo representations show the sequence motifs upstream the 998 primary TTSs and 1359 secondary TTSs.

**Figure 1- figure supplement 3.**
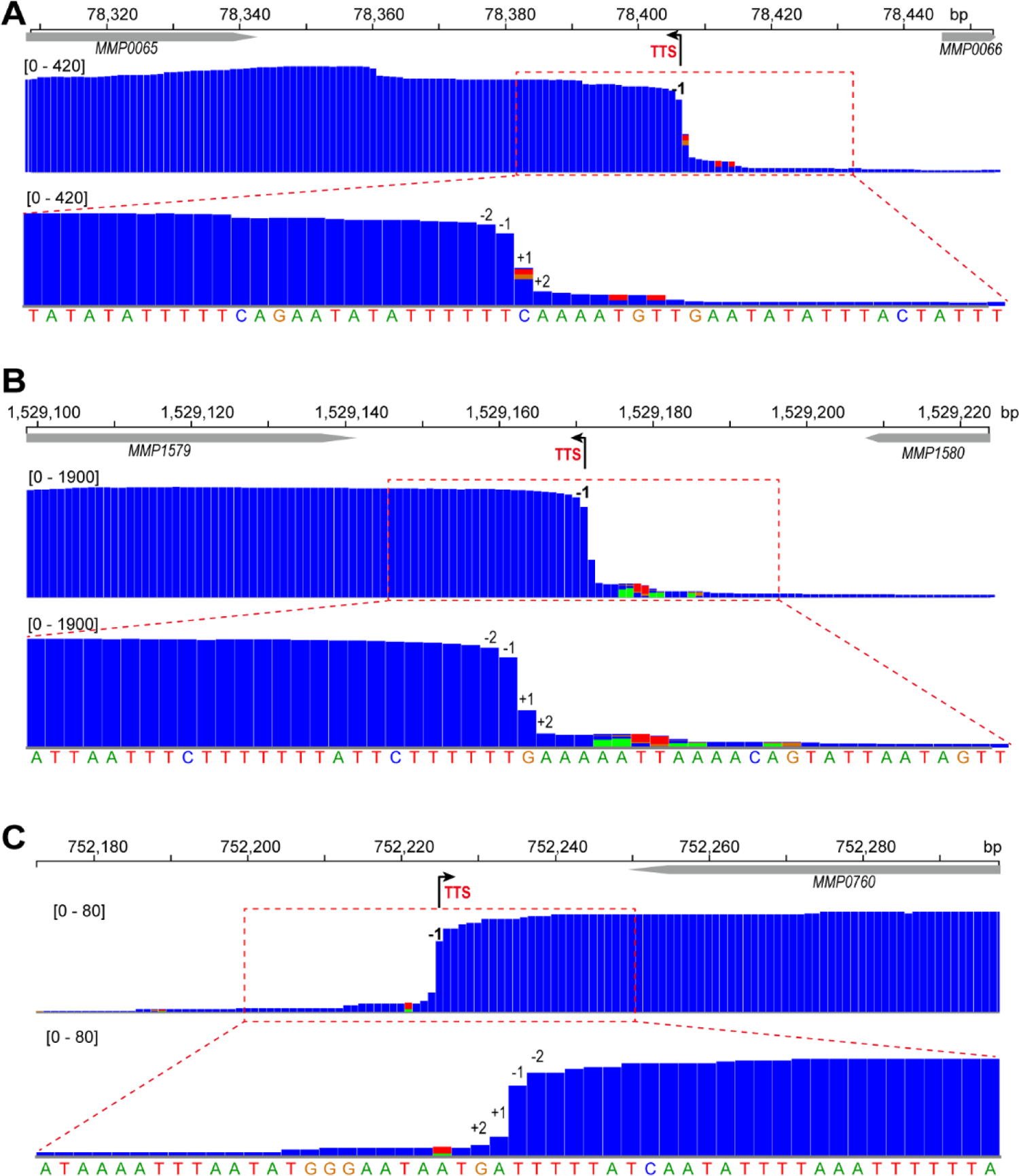
Term-seq mapping maps of three representative genes, *MMP0065* (A), *MMP1579* (B) and *MMP0760* (C). Gene locations at chromosome are indicated at the top. Black arrows indicate Term-seq identified transcription termination site (TTS, −1). The magnified mapping region (red dot framed) showed a step-wise reads decreasing pattern from −2 nt (upstream) to +2 nt (downstream) of TTS (lower panels). Numbers inside brackets are the Term-seq reads mapping heights.

**Figure 1- figure supplement 4.**
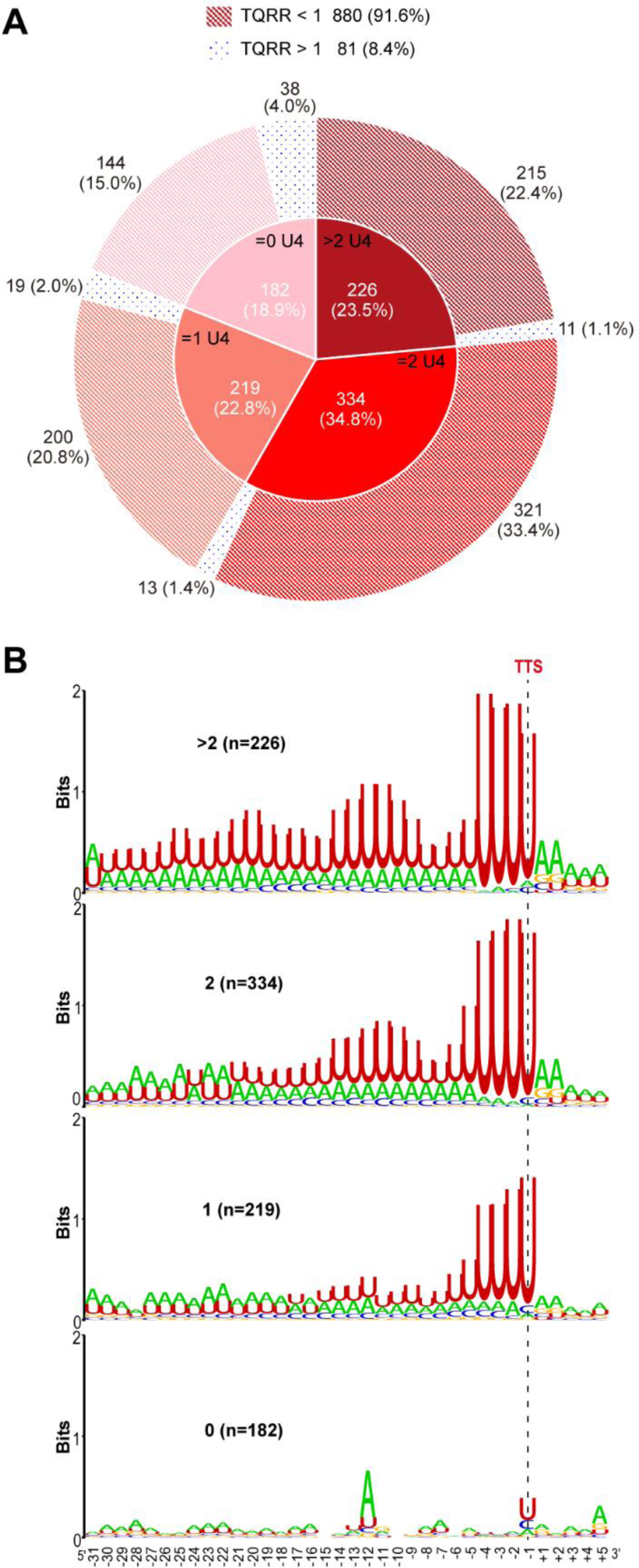
Percentages and motifs of the TU groups that have different terminator U-tracts. (A) The pie chart shows the TU numbers and percentages in each group that has >2, 2, 1, 0 U4-tracts, respectively. aCPSF1 dependency in each group with different terminator U-tract numbers determined by TQRR was displayed in the outer ring. (B) Logo representations of the terminator motif signatures that contain more than 2 (>2), 2, 1 and 0 U4-tracts, respectively. Inside parentheses are the terminator numbers that contain various U-tracts in each group.

Next, all of the identified TUs were ranked into three hierarchical groups according to the calculated TTEs: high TTE (>60%), medium TTE (30%< TTE <60%) and low TTE (<30%) groups. Statistically, about 32.5% and 44% of TUs fell in the high and medium TTE groups and only 23.5% of TUs fell in the low TTE group (Figure 1C). Sequence motifs generated from −30 nt until +5 nt flanking TTSs using Weblogo showed characteristic U-rich tracts, each containing four consecutive uridine nucleotides (U4) preceding TTSs among the overrepresented TUs in all the three groups (Figure 1C). Noticeably, a positive correlation appeared between TTE and the terminator U-tract length, such as two or more than two U4-tracts were found overrepresented in the high TTE group, while the U4-tract was underrepresented in the low TTE group (Figure 1C). To further evaluate the correlation between the U4-tracts and the TTEs, we classified all of the defined TUs into four groups based on the U4-tract numbers preceding the primary TTSs (Figure 1- figure supplement 4A), and then generated the sequence motif (Figure 1- figure supplement 4B) and statistically calculated the TTE distribution (Figure 1D) in each group. Similarly, a marked positive correlation between the U4-tract numbers and the TTEs was also observed, i.e., TUs with more U4-tract numbers having higher TTEs, such as TUs in the groups of >2, 2, 1 and 0 U4-tracts were found having the median TTEs of 55.5%, 52.1%, 43.3%, and 30%, respectively.

Therefore, the above analyses indicate that the U4-tract preceding the TTS could be a key signal (motif) in dictating or affecting RNAP to pause and in triggering transcription termination, and more U4-tracts could result in higher TTEs in *M. maripaludis*.

### Concurrence of the terminator U-tract and termination factor aCPSF1 in dictating high transcription termination efficiencies

Based on our recent finding that aCPSF1 functions as the archaeal general transcription termination factor in mediating genome-wide transcription termination and shaping the transcriptome of *M. maripaludis* (Yue et al., 2020), we quantitatively compared the TTEs in the wild-type and aCPSF1 expression depleted (▽*aCPSF1*) strains based on the Term-seq data, and found an average 50% reduction in TTEs of primary TTSs when aCPSF1 was depleted (Figure 2A and Figure 2- figure supplement 1). To further quantify the contribution of aCPSF1 to TTE, the aCPSF1 dependency of a TU in transcription termination was calculated based on its TTS quadruplet read changes in ▽*aCPSF1* compared to the wild-type strain using the following formula: TTS Quadruplet Read Ratio 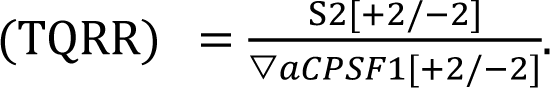 That TUs having TQRR <1, i.e., the read ratio between +2 and −2 nt in the TTS quadruplet is reduced due to aCPSF1 depletion, were identified as aCPSF1-dependent termination. Unexpectedly, we found 91.6% (880/961) TUs having TQRR <1 (Figure 1 - figure supplement 4A), and observed an approximately linear correlation between TQRR and TTE for all the studied 961 TUs (Figure 2B), indicating that majority of TUs were terminated in an aCPSF1-dependent manner, and the higher TTE of a TU, the more dependency of aCPSF1 in the TTE. Therefore, aCPSF1 could display a positive correlation with TTEs as well as the terminator U-tracts and play a key role in dictating high TTEs at the genome-wide level.

**Figure 2.**
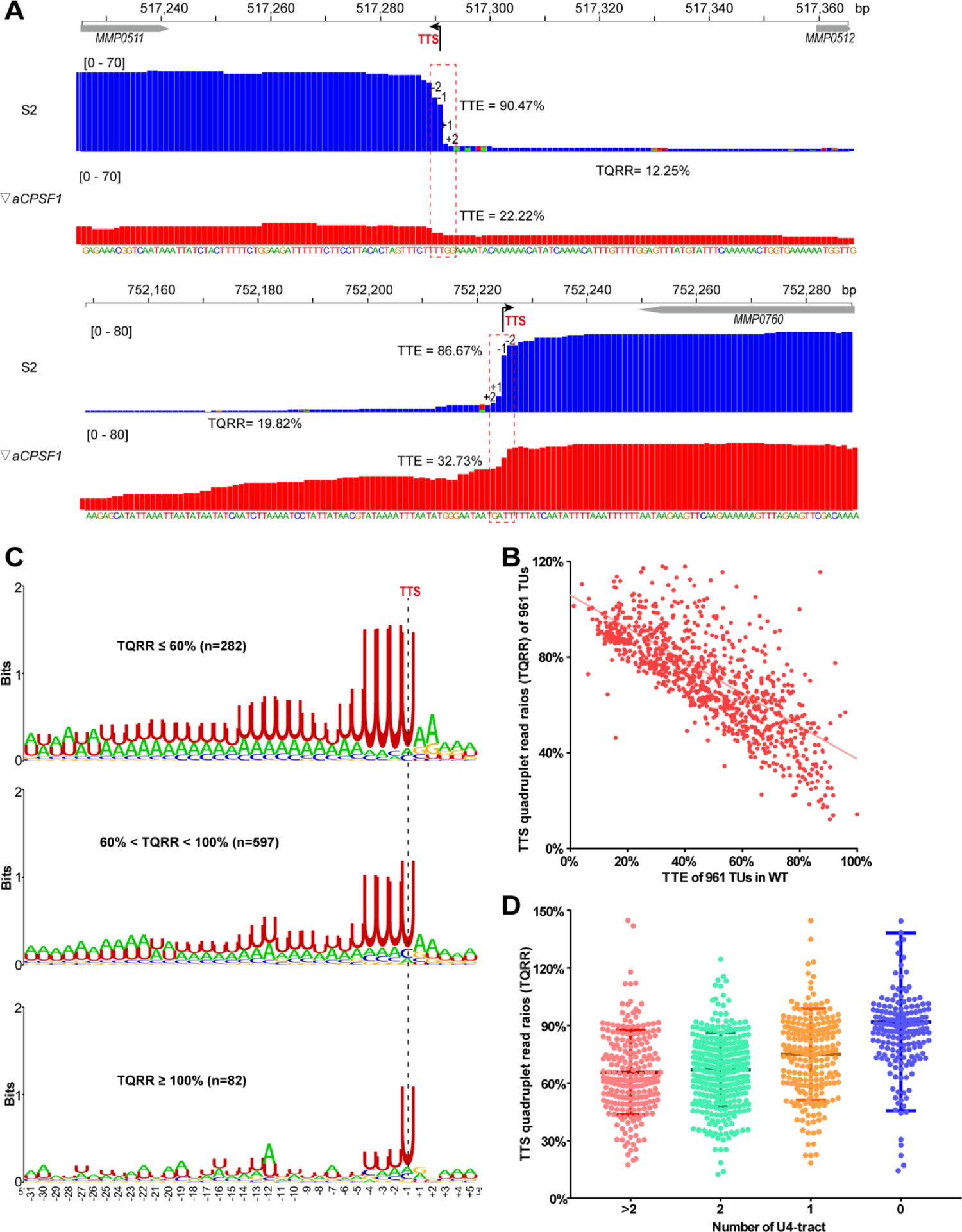
Co-occurrence of aCPSF1 and the terminator U4-tract is correlated to the genome-wide transcription termination efficiencies (TTEs) of *M. maripaludis*. *(A)* Visualized Term-seq read mapping maps of the representative genes, *MMP0511* (top) and *MMP0760* (bottom), show sharper reads decreasing between the −2 and +2 nts respectively down- and up-stream of the TTSs (dotted line) in the wild-type strain (S2) than in the aCPSF1 depletion mutant (▽*aCPSF1*). The chromosome locations of genes are indicated at the top. The bent arrow indicates the Term-seq identified transcription termination site (TTS, −1). TQRR represents the TTS quadruplet read ratio of a TU in the wild-type strain to that in the aCPSF1 depletion mutant, with the lower values showing a higher aCPSF1 dependency of a TU in transcription termination. Inside the brackets are the mapping read heights. TTE is calculated as above. *(B)* A linear correlation is observed between TQRR values and TTEs of 961 protein coding TUs. *(C)* Logo representations of the terminator motif signatures are shown for highly aCPSF1-dependent (TQRR≤60%), moderately aCPSF1-dependent (60%<TQRR<100%) and non-dependent (TQRR≥100%) groups. Inside the parentheses are the TU numbers of each group. *(D)* Box-plot diagrams showing the TQRR (aCPSF1 dependency) statistics of the terminators carrying >2, 2, 1, and 0 U4-tracts. The diagram representations are the same as in Figure 1B. The significance statistical analysis for the TQRRs of the four groups determined by Wilcoxon rank sum test were shown in Table S4.

**Figure 2- figure supplement 1.**
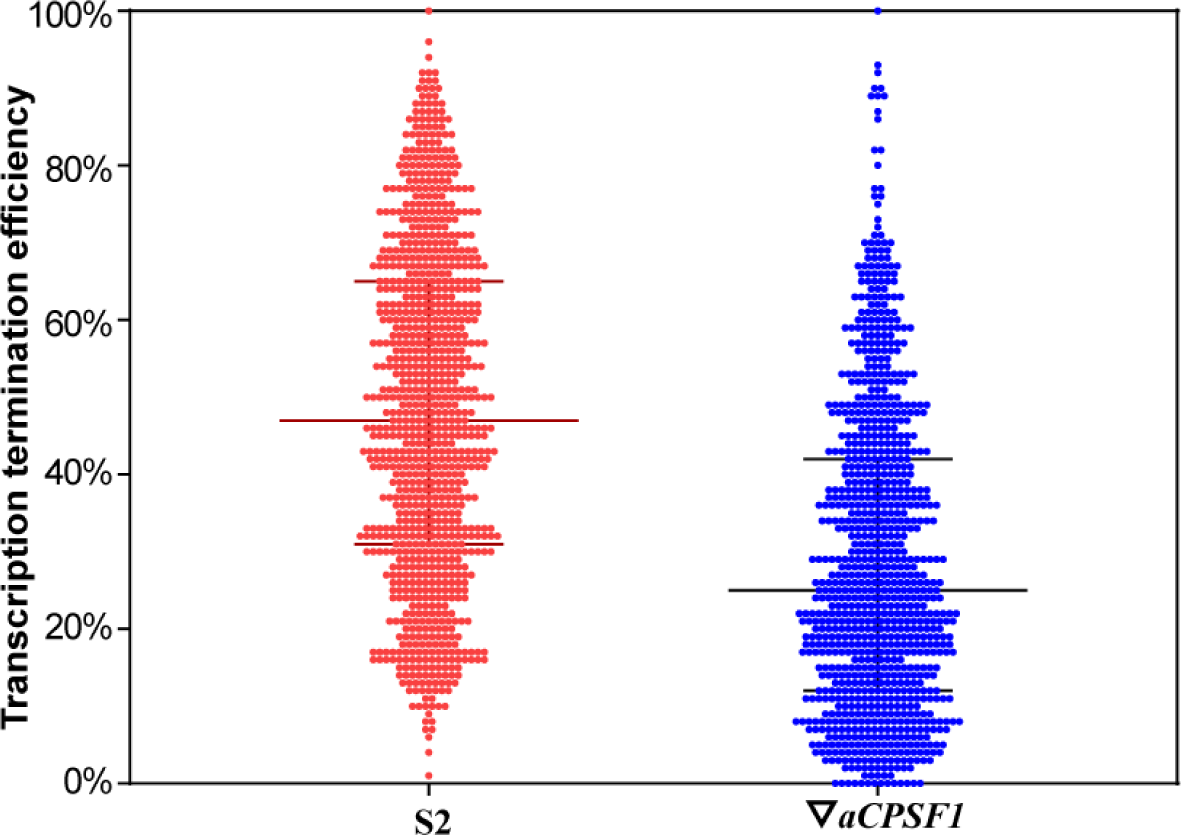
A box plot diagram shows the statistics of transcription termination efficiencies (TTEs) of the Term-seq identified primary TTSs in strain S2 and ▽*aCPSF1* mutant.

Subsequently, we explored the cis-elements that may determine the aCPSF1-dependent TTE, i.e., the cis-elements recognized by aCPSF1 by statistically analyzing the correlation between the aCPSF1 dependency of TTEs and the presence of sequence motifs preceding TTSs of the TUs. Based on TQRRs, we classified the above 961 TUs into three groups, i.e., the highly aCPSF1-dependent (TQRR≤60%), moderately aCPSF1-dependent (60%<TQRR<100%) and non-aCPSF1-dependent (TQRR≥100%) groups, and generated the TTS preceding sequence motif for each group using Weblogo. Interestingly, we not only found that 29.3% (282/961) and 62.2% (597/961) of TUs belonged to the highly and moderately aCPSF1-dependent groups, respectively, and only 8.4% (81/961) TUs felled into aCPSF1-nondependent group, but also found a significant positive correlation between the aCPSF1-dependency and the numbers of U4-tract preceding TTSs, i.e., TU groups with higher aCPSF1 dependency have more featured U4-tracts (Figure 2C). Additionally, we evaluated the relationship between the aCPSF1-dependency and the above four U4-tract TU groups analyzed in Figure 1D. Interestingly, we found that 94.5% (736/779) of TUs in the ≥1 U4-tract TU groups have a TQRR <1 (Figure 1 - figure supplement 4A), and also found that TU groups with ≥2, 2, 1 and 0 U4-tracts had the median TQRRs of 65.5%, 67%, 76%, and 92%, respectively, thereby indicating that majority of TUs with U4-tract depend on aCPSF1 for termination and the TU groups of containing more U4-tracts have lower median TQRR values, namely, higher aCPSF1-dependency (Figure 2D). Noteworthily, even among TUs with 0 U4-tract, 79.1% (144/182) of TUs were found having a TQRR <1 (Figure 1 - figure supplement 4A), suggesting that transcriptions of these TUs with weak terminator motif can also be warranted for termination by aCPSF1. Additionally, among the 8.4% (81/961) of TUs that felled into aCPSF1-nondependent group, only 3.95% (38 of 961) of TUs were found having 0 U4-tract (Figure 1 - figure supplement 4A), suggesting that these very few TUs may be terminated by mechanisms independent of both aCPSF1 and U4-tract terminator or the TTSs identified for these TUs could be potential RNA processing sites derived from stale RNA processing or degradation products.

Collectively, these results indicate that both the terminator cis-element U4-tracts and the trans-action termination factor aCPSF1 exhibit highly positive correlation with the TTEs, and the two appear synergistically dictating high TTEs at a genome-wide level *in vivo*; in addition, aCPSF1 likely backs up the termination of TUs without the terminator U4-tract.

### aCPSF1 specifically binds to RNAs embedding the terminator U4-tract sequence ***in vitro***

The collaboration of aCPSF1 and the terminator U-tract cis-element in dictating TTE suggests that this termination factor might specifically recognize the terminator U-tract motif embedded in the nascent transcript 3′-end to dictate archaeal transcription termination. To test this hypothesis, we then first assayed the binding ability of aCPSF1 to three synthetic RNAs that contain the terminator U-tract sequences of transcripts *MMP0901*, *MMP1149* and *MMP1100*, which were determined to be cleaved by the recombinant aCPSF1 in our previous study (Yue et al., 2020). An RNA fragment with the transcript 3′-end sequence of *MMP1697* lacking a U-tract was included as a control. Using RNA electrophoretic mobility shift assay (rEMSA), shifted protein–RNA complex bands could be observed in the three U-tract containing RNAs, but not in that without U-tract at the same aCPSF1 contents (Figure 3 - figure supplement 1). Next, 12 additional RNA fragments in a consensus 36 nt length and of transcript 3′-end sequences, from 30 nt upstream to 5 nt downstream from the TTS, were used to compare the binding ability of aCPSF1 to them. These sequences were derived from six transcripts embedding ≥2 U4-tracts (Figure 3A), three transcripts embedding 1 U4-tract (Figure 3B) and three with no (0) U4-tract (Figure 3C), respectively. The rEMSA results indicated that aCPSF1 exhibited strongest binding to those with ≥2 U4-tracts, weaker binding to those with one U4-tract and weakest binding to those without U-tract (Figure 3). Supportively, using surface plasmon resonance (SPR) assay and at the same contents of aCPSF1, the highest resonance unit (RU) was determined for the RNA derived from *MMP0400* 3′-end sequence with the longest U-tract, but the lowest for that from *MMP1406* 3′-end carrying the shortest U-tract (Figure 3 - figure supplement 2). Therefore, these results demonstrated that aCPSF1 specifically recognizes the transcripts with U-tract sequences and binds preferentially to the transcripts carrying more U4-tracts at the 3′-ends.

**Figure 3.**
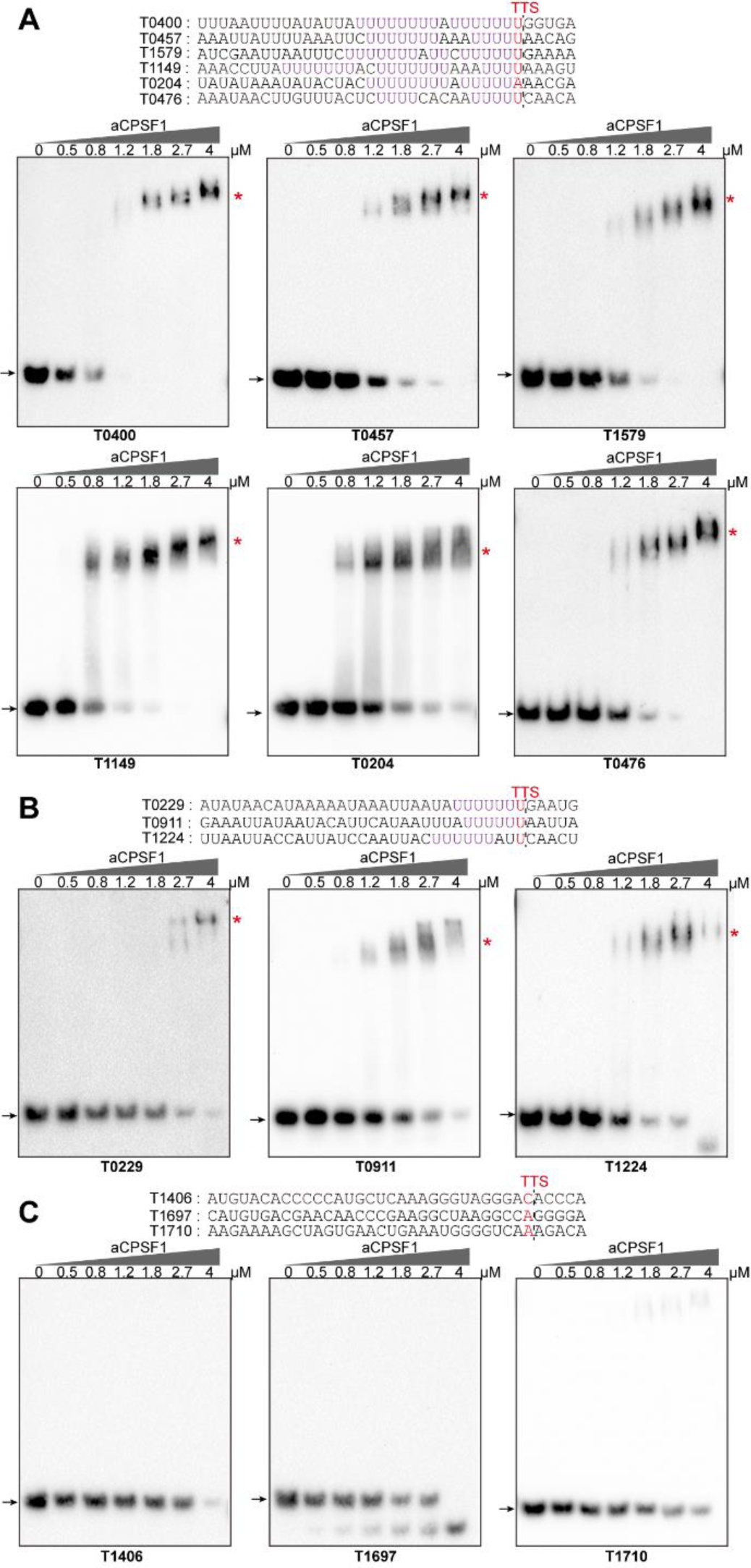
rEMSA assays the aCPSF1 binding specificity to RNAs carrying different numbers of U4-tracts. RNAs with a consensus length of 36 nt derived from the indicated gene terminators that carry two U-tracts *(A)*, one U-tract *(B)* and no U-tract *(C)* were used as the binding substrates. RNA sequences are shown in the top panels by red letters indicating Term-seq identified TTSs. The gradient concentrations of aCPSF1 used in the binding mixtures are indicated at the top of gels. Detailed binding procedure is described in the Methods section, and chemiluminescence signals were visualized on a Tanon-5200 Multi instrument. Arrows and red stars indicate the free RNA substrates and the RNA-aCPSF1 complexes, respectively. Binding assay for each RNA substrate was performed for triplicate measurements.

**Figure 3- figure supplement 1.**
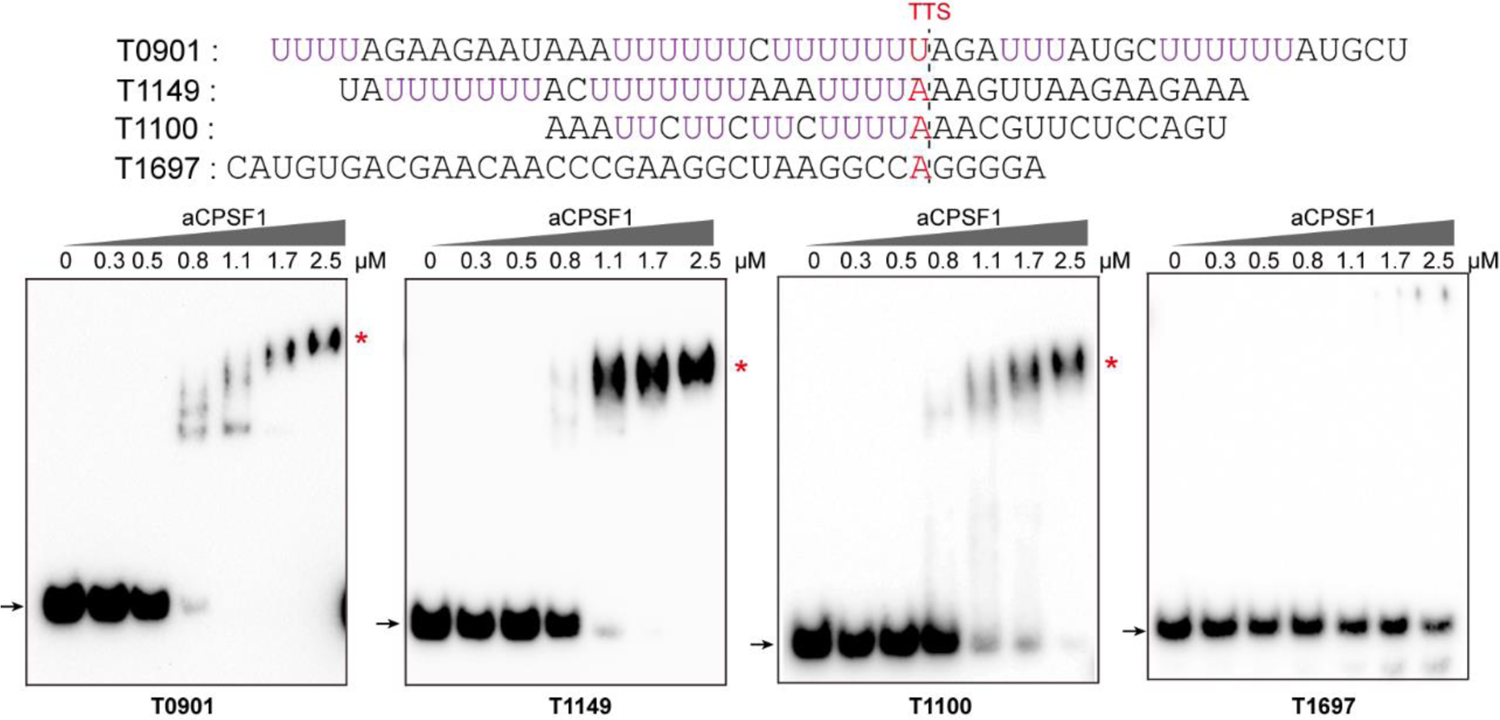
rEMSA assayed the U-tract binding specificity of aCPSF1. Three RNAs (T0901, T1149 and T1100) containing the terminator U-tract sequences and one without U-tract (T1697), that are from the indicated transcript 3′-ends, are used as aCPSF1 binding substrates. The terminator sequences are shown in upper panels, and U-tract sequences and Term-seq identified TTS are shown as magenta and red letters, respectively. The gradient concentrations of aCPSF1 used in the binding mixtures are indicated at the top of gels. Black arrows indicate free RNAs and red stars specify RNA-aCPSF1 complexes.

**Figure 3- figure supplement 2.**
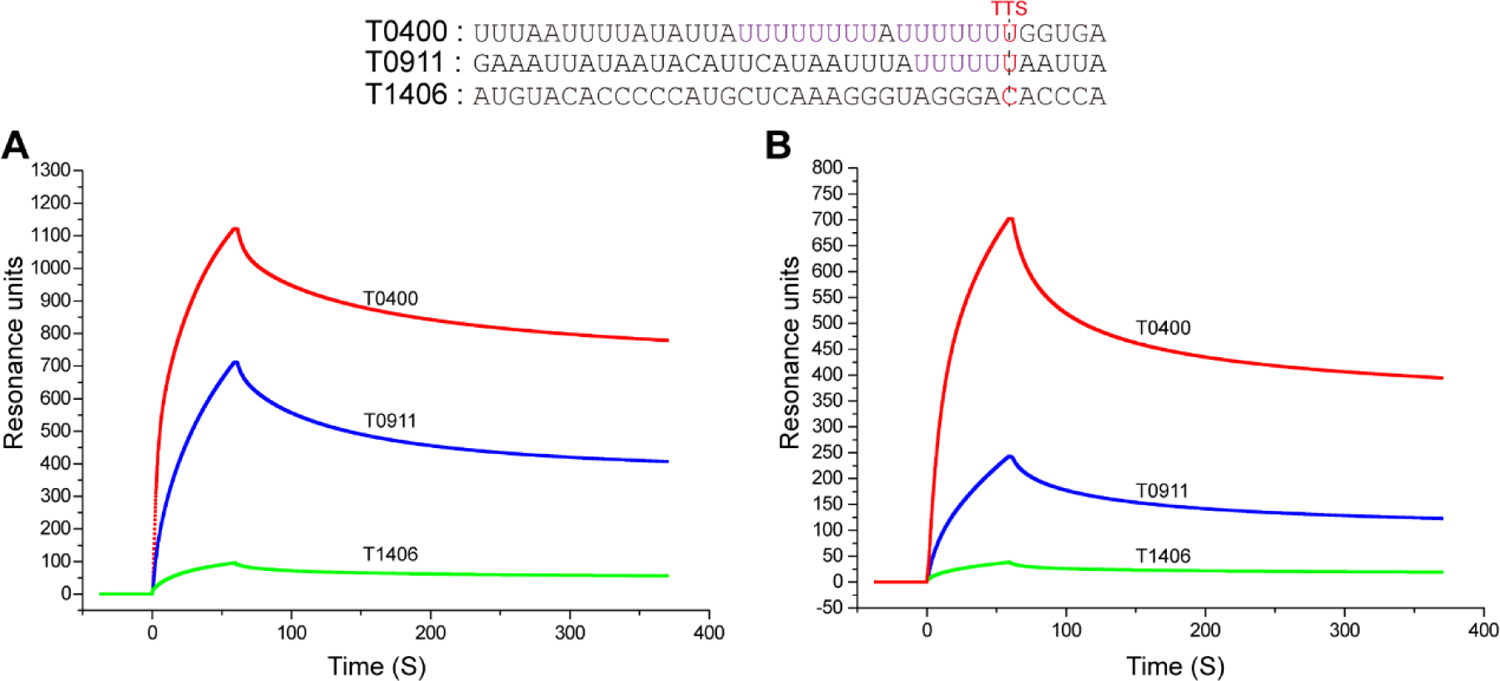
SPR assayed the U-tract binding specificity of aCPSF1. Two RNAs of terminator sequences carrying U-tracts (T0400 and T0911) and one without U-tract (T1406) are used as the binding substrates for SPR assays. RNA sequences are shown at the top. The purified recombinant *Mmp*-aCPSF1 at 500 µg/ml (A) and 125 µg/ml (B) were assayed for RNA binding.

Next, the minimal RNA length and U-tract sequence required for aCPSF1 binding were investigated. The 36 nt RNA sequences from the transcript 3′-ends of *MMP0204* and *MMP0400*, embedding ≥2 U4-tracts, were sequentially truncated by 6 nt from the 5′-end to generate 30 nt, 24 nt, and 18 nt RNA substrates. The rEMSA results indicated that aCPSF1 bound to the 36 nt, 30 nt and 24 nt RNAs with similar affinity, but had a sharply reduced affinity to the RNA of 18 nt (Figure 4A and Figure 4 - figure supplement 1A). To confirm the role of the U-tracts in determining the binding specificity of aCPSF1, base mutation was performed on either one U-tract (18nt-M1 and 18nt-M2) or both U-tracts (18nt-M3) of the 18 nt RNAs. The rEMSA results indicated that mutation of either one U-tract or both U-tracts remarkably reduced the binding ability of aCPSF1 to the RNA substrate (Figure 4B and Figure 4 - figure supplement 1B). Reciprocally, by mutating two As to Us on the RNA with the *MMP0229* 3′-end sequence, which originally has only one U-tract, to obtain two U-tracts on the mutated RNA of T0229-18nt-M1, the binding ability of aCPSF1 to T0229-18nt-M1 was notably improved compared to T0229-18nt, whereas mutation of the four Us to four As at T0229-18 nt to obtain an RNA sequence lacking U-tract (T0229-18nt-M2) completely abolished aCPSF1 binding (Figure 4C). Therefore, these results demonstrated that either one of the two U-tracts in the transcript 3′-end is necessary for efficient and specific binding of aCPSF1. Furthermore, the minimum length of the consecutive uridines in the double U-tracts was evaluated. The double U-tracts RNA of T0204-24nt was used as the template for U to C mutation, generating T0204-DU5, T0204-DU4 and T0204-DU3, which carry 5, 4, or 3 consecutive uridines in each U-tract, respectively. The rEMSA results indicated that aCPSF1 efficiently bound to T0204-DU5 and T0204-DU4, but could not bind to T0204-DU3, thus suggesting that the double U4-tracts could be the minimum terminator sequence for efficient binding of aCPSF1 (Figure 4D). Collectively, the *in vitro* RNA binding experiments demonstrated that an RNA fragment embedding a double U4-tract and with a length of at least 18 nt is the cis-element required by the termination factor aCPSF1 for efficient and specific binding.

**Figure 4.**
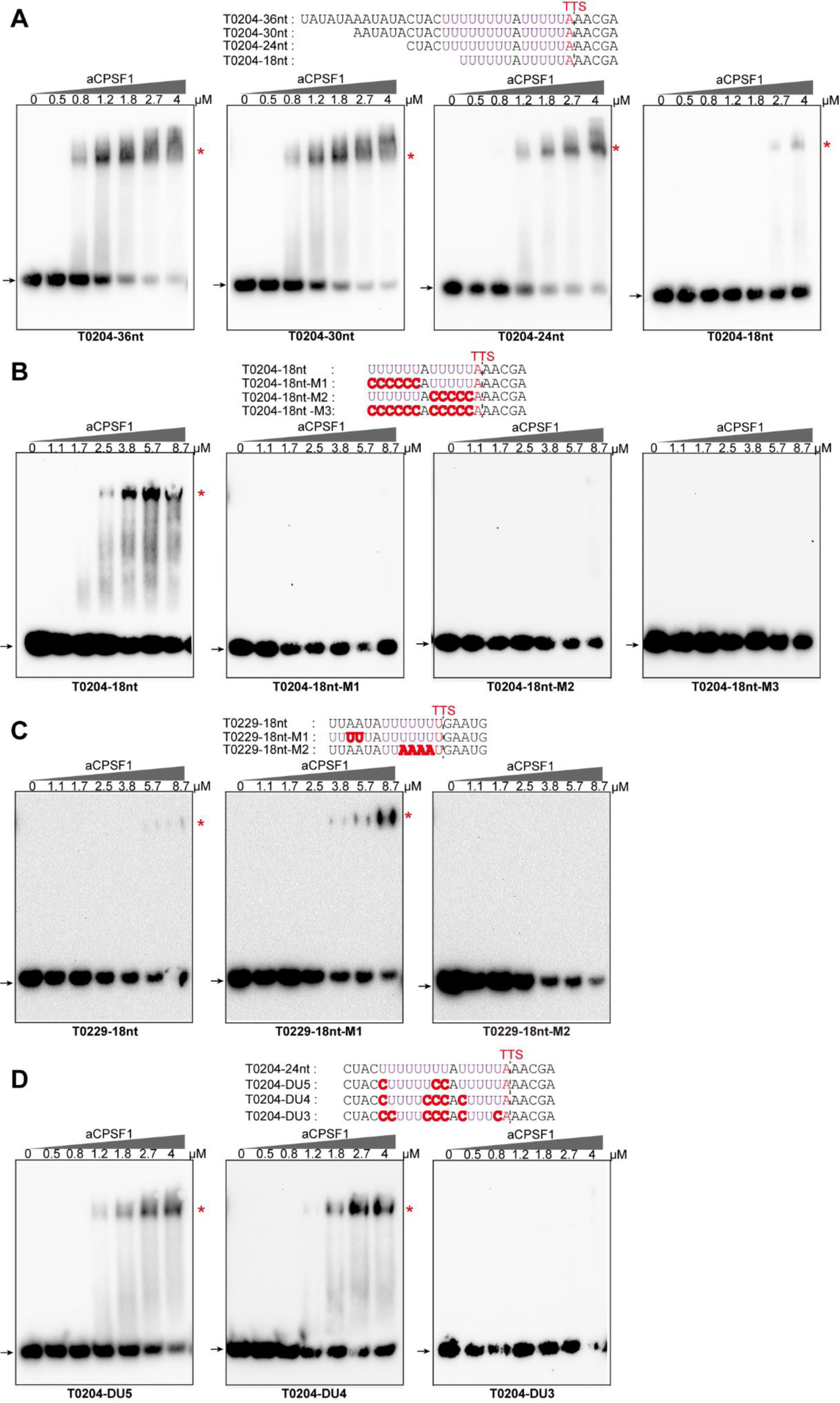
rEMSA assays the minimal RNA length and U-tract base stringency **required by aCPSF1.** RNAs of indicated lengths and base mutations shown in the top panels derived from the native terminator sequences of *MMP0204* and *MMP0229* were used as the binding substrates of aCPSF1. *(A)* The *MMP0204* terminator RNA with a length of 36 nt was truncated by 6 nt from the 5′ end, resulting in RNAs with lengths of 30 nt, 24 nt, and 18 nt. *(B)* The 18nt-*MMP0204* terminator RNA was base mutated in either one (M1 and M2) or both U-tracts (M3). *(C)* A one U-tract RNA derived from the *MMP0229* terminator was base mutated to construct a two U-tract mutant (M1) and a non-U-tract mutant (M2). *(D)* The two U-tract contained RNA from the 3′-end of *MMP0204* was base mutated to generate the two U-tracts with each U-tract containing 5 (DU5), 4 (DU4), and 3 (DU3) consecutive Us, respectively. TTSs identified by Term-seq and the mutated residues are shown as plain and bold red letters, respectively. The gradient concentrations of aCPSF1 used in the binding mixtures are indicated at the top of gels. Detailed binding procedure is described in the Methods section, and chemiluminescence signals were visualized on a Tanon-5200 Multi instrument. Arrows and red stars indicate the free RNA substrates and the RNA-aCPSF1 complexes, respectively. Binding assay for each RNA substrate was performed for triplicate measurements.

**Figure 4- figure supplement 1.**
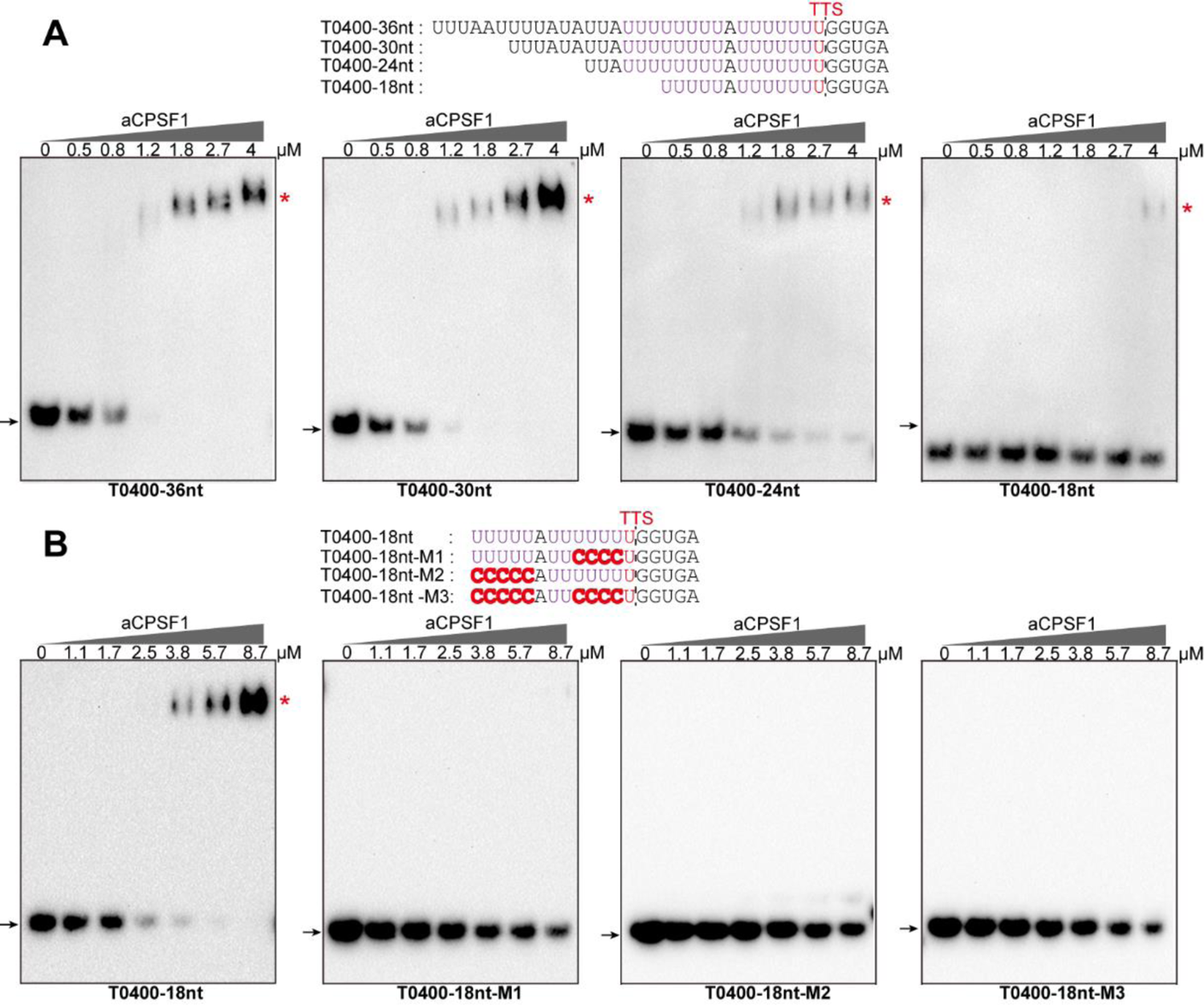
rEMSA assays the minimal RNA length and U-tract base stringency required by aCPSF1. RNAs in indicated lengths and base mutations shown at the top panels that are derived from the native terminator sequences of *MMP0400* were used as the binding substrates of aCPSF1. (A) The *MMP0400* terminator RNA in length of 36 nt was each 6 nt truncated from the 5′ end to result RNAs in lengths of 30 nt, 24 nt and 18 nt, respectively. (B) The 18nt-*MMP0400* terminator RNA was base mutated on either one (M1 and M2) or two U-tracts (M3). TTSs identified by Term-seq and the mutated residues are shown as plain and bold red letters, respectively. The gradient concentrations of aCPSF1 used in the binding mixtures are indicated at the top of gels. Detailed binding procedure was described in the Methods section, and chemiluminescence signals were visualized on a Tanon-5200 Multi instrument. Arrows and red stars indicate free RNA substrates and RNA-aCPSF1 complexes, respectively.

### An *in vivo* synergism of aCPSF1 and the terminator U-tract dictates the effective transcription termination

To verify the *in vivo* synergistic action of aCPSF1 and the terminator U-tract in dictating TTEs, indispensability of the two for efficient transcription termination was assayed. A dual reporter gene system was constructed as shown in Figure 5A, in which the tested terminators, in length of 36 nt with the same sequences as in the rEMSA assays (Figure 3), were each inserted between the genes encoding luciferase and mCherry fluorescent proteins. In total, five terminator reporters were constructed, among which T1149 and T0204 carry two U-tracts, T0229 and T0911 carry one and T1710 carries none. The reporter constructs were respectively transformed into the wild-type strain S2, an aCPSF1 depletion mutant (▽*aCPSF1*), and two complementary strains, Com(WT) and Com(Mu), which carry the wild-type aCPSF1 and its catalytically inactive mutant complementation in ▽*aCPSF1*, respectively. TTE was assayed based on the transcript abundance (TA) of the downstream mCherry and the upstream luciferase, and calculated using the following formula: 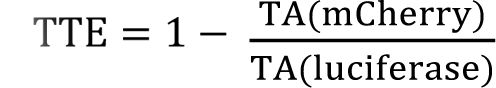. The attempt to using fluorescence to indicate transcript levels failed due to intensity fluctuations during the assay. Using quantitative reverse-transcription (RT-qPCR), transcript abundances of the two reporter genes were assayed, which determined that TTEs were highest for the terminators with two U-tracts (T1149 and T0204), lower for those with one U4-tract (T0911 and T0229), and the lowest for that without U-tract (T1710) in the wild-type strain S2 (Figure 5B). Therefore, the reporter gene system confirms the positive correlation between the number of terminator U-tracts and TTEs *in vivo*.

**Figure 5.**
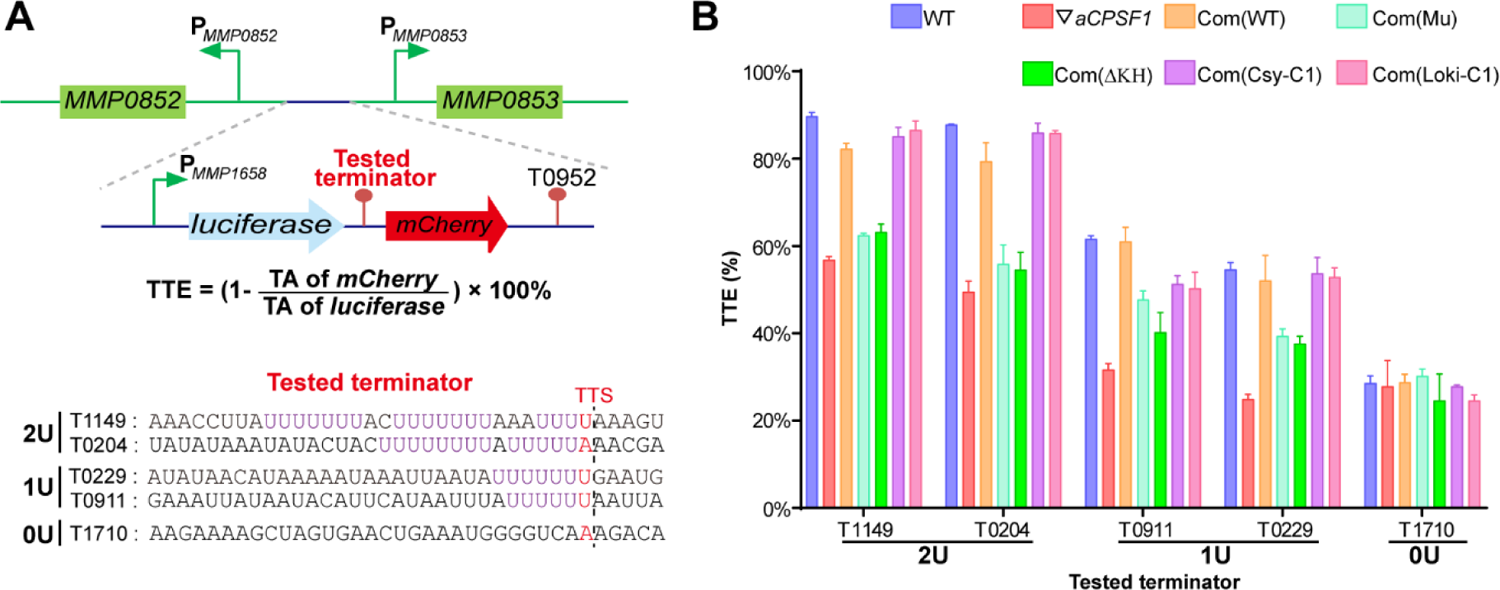
A terminator reporter system assays TTE variations when aCPSF1 and the terminator U-tract co-occur or absence of either one. (*A*) A schematic depicts the construction of the terminator reporter system. The tested terminator sequences carrying different numbers of U-tracts were each inserted between the upstream luciferase and downstream mCherry genes, and then fused downstream of the *MMP1658* promoter (P_MMP1658_) and upstream of the *MMP0952* terminator (T0952). Subsequently, the constructed DNA fragment was inserted in between the promoters of *MMP0852* and *MMP0853* in the genomes of various strains listed in (*B*) through homologous recombination. Detailed protocol is described in the Methods section. The formula for TTE calculation and the sequences of tested terminators are shown below. *(B)* Quantitative RT-PCR analysis was conducted to quantify the luciferase and mCherry transcript abundances in various terminator constructs that were expressed in the wild-type strain (WT), the aCPSF1 depletion mutant (▽*aCPSF1*), and the ▽*aCPSF1* strain complemented with wild-type (Com(WT)), KH domain truncated (Com (ΔKH)) and catalytic mutated (Com(Mu)) of *M. maripaludis* aCPSF1 (*Mmp*-aCPSF1), and the aCPSF1 orthologs from Ca. *Lokiarchaeum* sp. GC14_75 Com (*Loki*-aCPSF1)and Ca. *Cenarchaeum symbiosum* Com(*Csy*-aCPSF1). TTEs were calculated based on the formula in the middle panel of (*A*), and TA indicates transcription abundance. Triplicated cultures were assayed and the averages and the standard derivations are shown. 2U, 1U, and 0U indicate the tested terminators carrying two (T1149, T0204), one (T0911, T0229), and 0 (T1710) U4-tracts, respectively. The statistical significance for the qRT-PCR analysis in different genetic strains Vs. WT determined by T-test were shown in Table S5.

However, in the▽*aCPSF1* strain, terminators containing either two U4-tracts (T1149 and T0204) or one U4-tract (T0911 and T0229) all exhibited reduced TTEs, while the terminator that contains no U-tract (T1710) retained nearly the same TTE as that in the wild-type strain (Figure 5B). Moreover, we calculated the aCPSF1 dependency in the dual reporter gene assays using the formula: 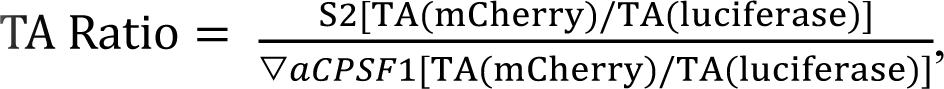 which is very similar to that for TQRR calculation.

TA ratios of 24%, 24%, 56%, 61%, and 99% were determined for the terminators carrying two U4-tracts (T1149 and T0204), one U4-tract (T0911 and T0229), and no U-tract (T1710), respectively. This result further verifies that terminators carrying more U4-tracts have a higher aCPSF1-dependency. It is worth noting that similar reduced TTEs were determined for the U-tract containing terminators in the Com(Mu) strain as those in ▽*aCPSF1*, while the TTEs in the Com(WT) strain were similar to those in the wild-type strain (Figure 5B). This indicates that the nuclease activity of aCPSF1 is also essential to the TTE dictation of the U-tract containing terminators. Thus, the dual reporter gene assays verified the *in vivo* synergism of the termination factor aCPSF1 with the terminator U-tract in dictating effective TTE in *M. maripaludis*, and the nuclease activity of aCSPF1 is necessary for the TTE dictation.

### N-terminal KH domains equip aCPSF1 to specifically bind the terminator U-tract and dictate transcription termination efficiency

The resolved crystal structure of an aCPSF1 ortholog from *Methanothermobacter thermautotrophicus* has revealed a tripartite architecture consisting of two N-terminal KH domains (KHa and KHb), a central MβL domain, and a C-terminal β-CASP domain (Silva et al., 2011). KH domains have been predicted to be involved in RNA sequence recognition and binding (Phung et al., 2013; Silva et al., 2011). Therefore, to assess whether the KH domains equip aCPSF1 to recognize the terminator U-tract, the N-terminal fragment of *Mmp*-aCPSF1 in length of 149 amino acids that contains the two KH domains was expressed, and the purified KH protein was assayed for the binding abilities to the RNAs and compared to the whole-length aCPSF1. As the intact aCPSF1 protein, the recombinant KH domains exhibited higher binding abilities to the RNAs containing longer or more U-tracts (T0204, T0400, T0457, T1579 and T0229) than to those without U-tract (T1406 and T1697) (Figure 6A). Therefore, the N-terminal KH domains enable aCPSF1 to specifically recognize the consecutive U-tracts embedded in the transcript 3′-ends.

**Figure 6.**
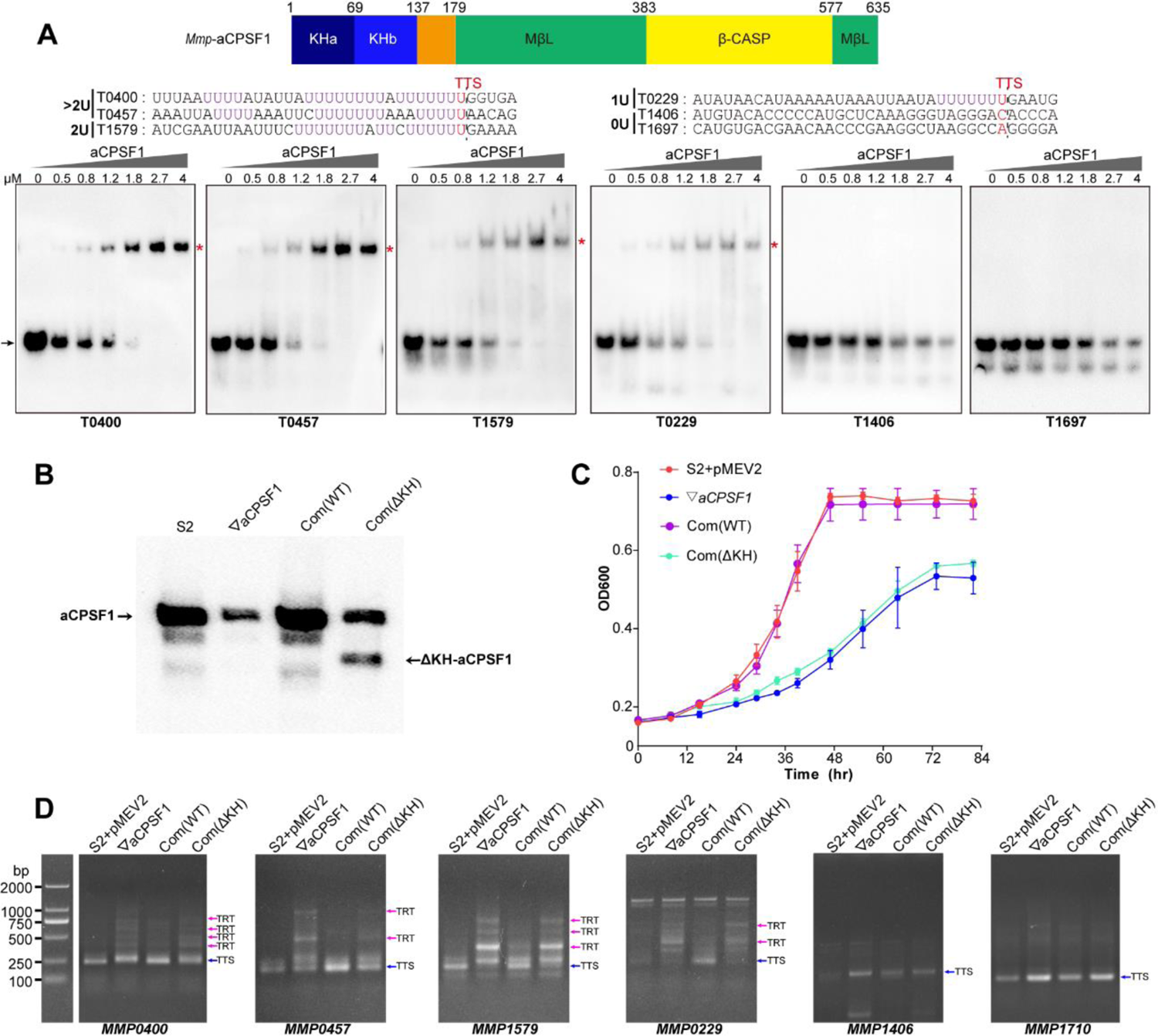
N-terminal KH domains of aCPSF1 contribute to the binding specificity to terminator U-tracts and transcription termination. *(A)* A schematic (upper panel) shows the aCPSF1 protein architecture with two N-terminal KH domains, the central MβL domain and the C-terminal β-CASP domain. rEMSA (lower panel) was performed to assay the binding specificity of the KH domains to the terminators carrying different numbers of U4-tracts as indicated. >2U, 2U, 1U, and 0U indicate the tested RNAs carried two, one, and no U4-tracts, respectively. The assayed RNA sequences are shown at the top of gels with Term-seq identified TTSs shown in red. rEMSA was performed as in Figure 3, and the added gradient concentrations of aCPSF1 KH domain are indicated. Arrows indicate free RNAs and red stars indicate the RNA–aCPSF1 complexes. *(B)* Western blot assays the expressions of the intact and KH domain deleted aCPSF1 proteins in the wild-type strain S2 carrying the empty complementation plasmid of pMEV2 (S2+pMEV2), the aCPSF1 depletion mutant (▽*aCPSF1*), and the ▽*aCPSF1* strain complemented with the wild-type (Com(WT)), and KH domain truncated (Com (ΔKH)) *Mmp*-aCPSF1. Arrows indicate the respective proteins. *(C)* Growth of the four strains was assayed on three batches of 22℃-grown cultures, and the averages and standard deviations are shown. *(D)* 3′RACE assays the transcriptional readthroughs (TRTs) in 22℃-grown strains with the same symbols as in (*B*). Blue and magenta arrows indicate the PCR products of normal terminations (TTSs) and TRTs, respectively. M, A DNA ladder is shown on the left (lane M).

**Figure 6- figure supplement 1.**
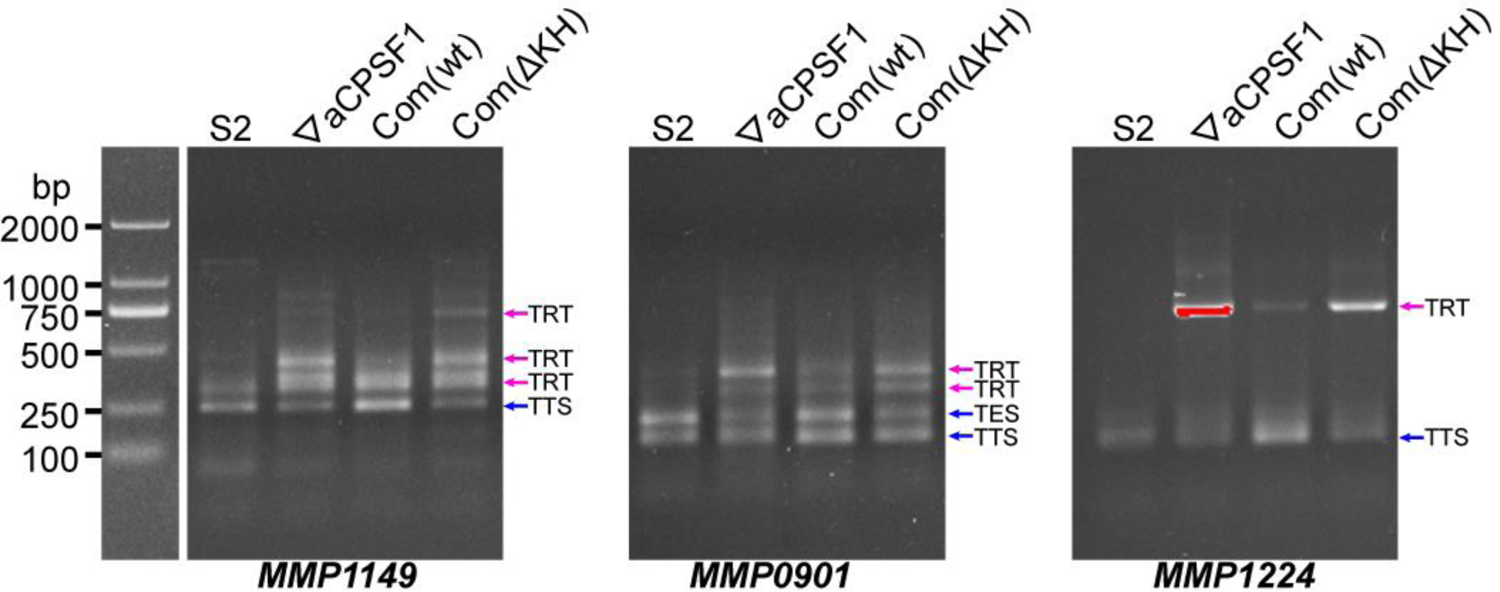
3′RACE assays transcriptional readthroughs (TRTs) of *MMP1149*, *MMP0901* and *MMP1224* in 22°C-grown wild type strain (S2), *aCPSF1* depletion mutant (▽*aCPSF1*), complementation of the wild-type (Com(WT)), and KH domain truncated (Com (ΔKH)) aCPSF1 at the background of ▽*aCPSF1*. Blue and magenta arrows indicate the PCR products of normal terminations (TTSs) and TRTs, respectively. A DNA ladder at left provides references of the PCR product migrations.

To further evaluate the role of the aCPSF1 KH domains in *in vivo* transcription termination, we complemented into ▽*aCPSF1* with the *aCPSF1* gene lacking the KH domains to obtain the complementary strain Com(ΔKH). The expression of the truncated aCPSF1 protein (ΔKH-aCPSF1) in strain Com(ΔKH) was assayed by western blot (Figure 6B). However, compared with strain Com(WT), which was complemented with the whole-length wild-type *aCPSF1* gene and restored the growth defect of ▽*aCPSF1*, strain Com(ΔKH) exhibited the same growth defect as ▽*aCPSF1* (Figure 6C). Furthermore, 3′RACE assays also detected similar transcription readthrough (TRT, transcription termination defect) products in seven transcripts with ≥1 U-tract embedded at the 3′-ends (*MMP0901*, *MMP1149*, *MMP0400*, *MMP0457*, *MMP1579*, *MMP0229* and *MMP1224*) in strain Com(ΔKH) as that in ▽*aCPSF1*, while no TRT products were detected in these seven transcripts in strain Com(WT) (Figure 6D and Figure 6 - figure supplement 1). In contrast, either in strain Com(ΔKH) or ▽*aCPSF1*, no TRTs were found in the transcripts without 3′-end U-tracts, *MMP1710* and *MMP1406* (Figure 6D). Together, these results suggested that the KH domains are essential for aCPSF1 to perform regular transcription termination of the transcripts carrying U-tract terminators and maintain the normal growth of *M. maripaludis*.

To further evaluate the contributions of the KH domains to aCPSF1-dependent TTEs in the transcripts containing terminator U-tracts *in vivo*, the five terminator reporters constructed above were transformed into strain Com(ΔKH). The TTEs of these reporters in Com(ΔKH) were determined similarly as above using the formula: 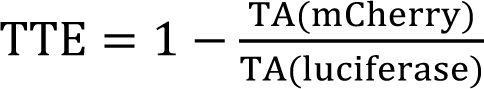. Transcript abundance quantification for the downstream mCherry and upstream luciferase genes showed that in contrast to the strain Com(WT) that was complemented with wild-type *aCPSF1*, complementation with *aCPSF1* lacking KH domains could not restore the TTE reductions of all the tested transcripts containing U-tract terminators (Figure 5B, Com(ΔKH)). Therefore, combination of the *in vitro* and *in vivo* experiments determined that the N-terminal KH domains are necessary for aCPSF1 to specifically recognize and collaborate with the terminator U-tracts in dictating effective TTEs.

### Synergism of aCPSF1 and the terminator U-tract in dictating transcription termination efficiency can be widely employed in Archaea

In Archaea, the U-rich terminators with 5, 6 and 7 consecutive Us (U5, U6, and U7) have been reported to be widely distributed in transcript 3′-ends (Berkemer et al., 2020; Dar et al., 2016a; Maier and Marchfelder, 2019; Yue et al., 2020); meanwhile, the aCPSF1 orthologs are also found to be strictly conserved and appear being vertically inherited (Phung et al., 2013; Yue et al., 2020). Thus, we asked whether the synergistic mode of aCPSF1 and the terminator U-tract is a common termination mechanism in Archaea. To test this hypothesis, two aCPSF1 orthologs, *Loki*-aCPSF1, from Ca. *Lokiarchaeum* sp. GC14_75 belonging to Lokiarchaeota, and *Csy*-aCPSF1, from Ca. *Cenarchaeum symbiosum* affiliating with Thaumarchaeota respectively, were chosen, as the two were determined capable of implementing the transcription termination function in *M. maripaludis* though they exhibit only 48% and 40% amino acid sequence identities with *Mmp*-aCPSF1 (Yue et al., 2020). The above five tested terminator reporters were transformed into strains Com*(Loki*-aCPSF1) and Com (*Csy*-aCPSF1), which were generated by complementing into strain ▽*aCPSF1* with the *Loki*-aCPSF1 and *Csy*-aCPSF1, respectively (Yue et al., 2020). Through quantifying transcript abundances of the reporter genes, TTEs of the tested terminators in Com*(Loki*-aCPSF1) and Com (*Csy*-aCPSF1) were calculated as described above. Noticeably, in strains Com*(Loki*-aCPSF1) and Com (*Csy*-aCPSF1), a similar U4-tract number related TTE pattern was determined as that in strain Com(WT), which was generated by complementing the wild-type methanococcal aCPSF1 into ▽*aCPSF1* as well (Figure 5B, Com(Loki-C1), Com(Csy-C1), and Com(WT)), i.e., the highest TTEs were determined for terminators with 2 U4-tracts (T1149 and T0204), and the TTEs with 1 U4-tract (T0911 and T0229) were lower. This result demonstrated that the heterogeneous aCPSF1 orthologs, although from distantly related archaea, could dictate the TTEs in *M. maripaludis* via distinguishing the terminator U-tracts as well. Given the ubiquitous distributions of both the termination factor aCPSF1 and the cis-element terminator U-tracts, the synergistic action mode of the two in dictating TTEs is predicted to be widely employed in various archaeal phyla.

## Discussion

Transcription termination controls the transcript boundaries and programmed transcriptome, and is hence one fundamental biological process (Porrua et al., 2016; Ray-Soni et al., 2016). However, our understanding of archaeal transcription termination mechanisms is still limited. Termination at the intrinsic terminator of a 3′-end U-tract cis-element was the sole known mechanism for a long time until most recently the protein factor aCPSF1 (FttA) was reported to achieve the genome-wide transcription termination and guarantee the transcriptome and cellular fitness (Sanders et al., 2020; Yue et al., 2020). However, aCPSF1-dependent termination was predicted to be only a back-up mechanism for the genes/operons that contain less efficient intrinsic termination signals and independent of the U-tract terminator mediated intrinsic termination (Sanders et al., 2020; Wenck and Santangelo, 2020). In this study, through a systematic analysis of the transcript 3′-end sequencing data in the model archaeon *M. maripaludis*, aCPSF1 was not only evidently found to function as a back-up mechanism to guarantee the terminations of TUs having weak intrinsic termination signal, but it was unexpectedly found a positive correlation with the genome-wide *in vivo* TTEs in concert with the terminator U-tracts numbers, indicating both the factor aCPSF1 and the intrinsic terminator U-tract cis-element collaboratively dictate the *in vivo* TTEs at the genome-wide level. By combining intensive *in vitro* and *in vivo* biochemical, genetic and molecular assays, this work has further elucidated that aCPSF1, through its N-terminal KH domains, specifically and hierarchically binds to the terminator U-tract signals embedded in transcript 3′-ends, and synergistically functions with this cis-element to elaborately dictate the effective genome-wide TTEs in *M. maripaludis*.

Therefore, distinct to the two independent termination mechanisms in bacteria, the factor Rho-dependent and intrinsic terminator dependent modes, the two-in-one termination mode based on the elaborate synergism of the factor aCPSF1 and the intrinsic terminator cis-element to control the *in vivo* high TTEs in Archaea is noteworthy, which could be the principal transcription termination mechanism in archaea. The transcript 3′-end cleavage triggered termination mechanism of aCPSF1 resembles the eukaryotic RNAP II termination mode (Yue et al., 2020). However, unlike eukaryotic RNAP II termination, where multiple protein factors are involved (Figure 7 and (Dominski, 2010; Eaton et al., 2018; Hill et al., 2019; Kuehner et al., 2011; Porrua et al., 2016)), the working mode of archaeal termination is also noteworthy as aCPSF1 may trigger termination by itself, using the N-terminal KH domain to specifically recognize and bind to the terminator U-tract and its β-CASP domain to implement transcript 3′-end cleavage, as depicted in Figure 7. Given the wide distribution of the terminator U-tract as well as the aCPSF1 orthologs among Archaea, and in view of the fact that two aCPSF1 orthologs of distant relatives exhibited the same synergism with the terminator U4-tracts, the two-in-one termination mode based on the synergism of the factor aCPSF1 and the terminator U-tracts in dictating high TTEs could be widely employed by archaeal phyla.

**Figure 7.**
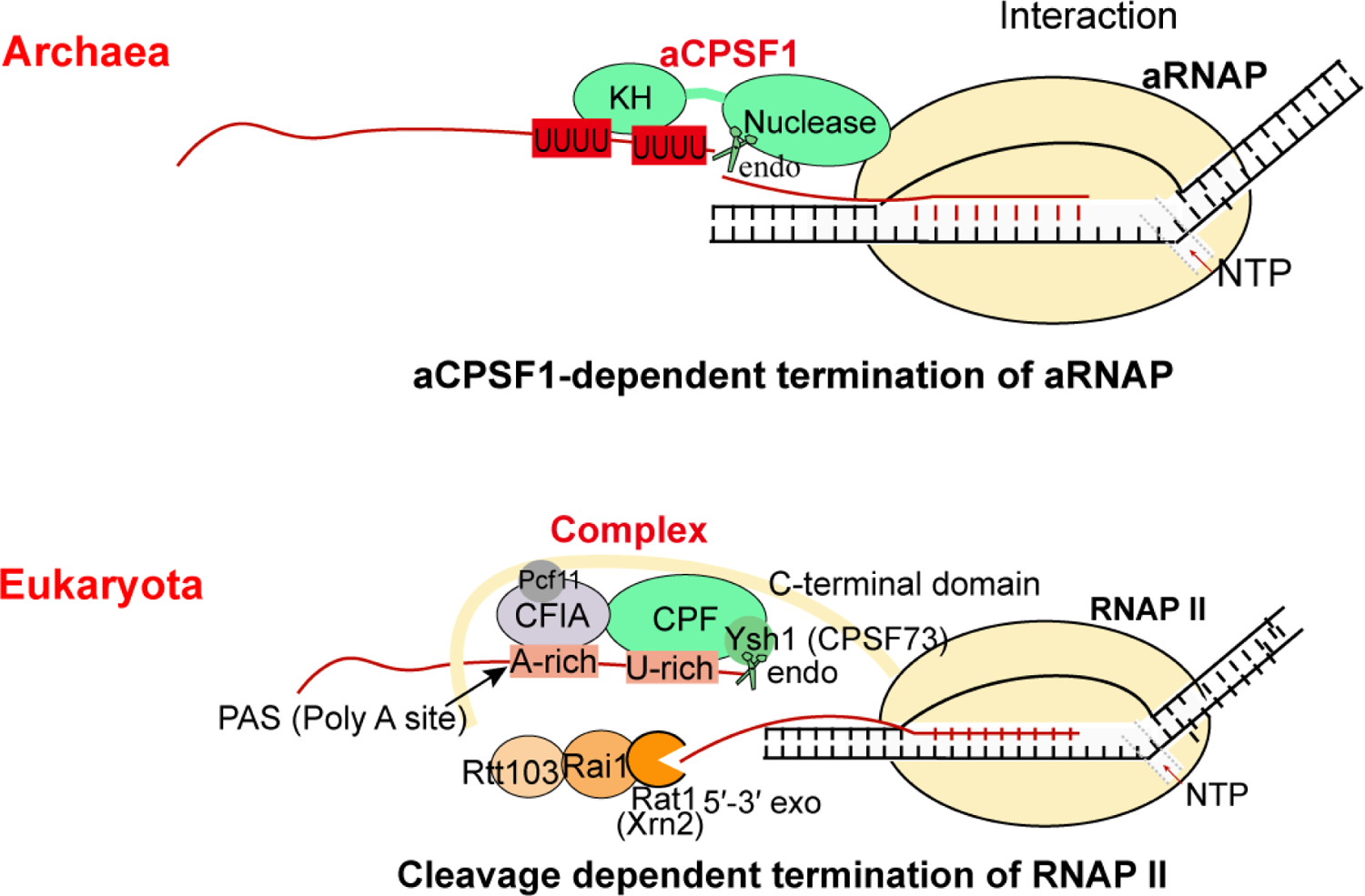
The aCPSF1-dependent archaeal transcription termination represents the archetype of the eukaryotic RNAP II termination mode. The general archaeal transcription termination factor aCPSF1, relying on the N-terminal KH domains specifically recognizing the terminator U-tract and the nuclease domain cleaving at the 3′-end, triggers transcription termination of archaeal RNA polymerase (aRNAP) (upper panel). The representative yeast RNAP II transcription termination model (Kuehner et al., 2011; Porrua et al., 2016; Porrua and Libri, 2015) is shown in the lower panel.

### aCPSF1 collaborates with the U-tract terminator signal to dictate archaeal transcription termination by specifically binding to it

Earlier studies on archaeal transcription termination have identified oligo(dT) stretches in non-template DNA strands usually upstream the TTSs of the coding and non-coding RNAs (Brown et al., 1989); therefore, the TTS proximal U-rich sequences were assumed to be the intrinsic terminator signal that triggers intrinsic termination events of archaeal RNAP analog to the bacterial one, which depends on a similar U-rich stretch but having a preceding hairpin structure (Brown et al., 1989; Maier and Marchfelder, 2019; Thomm et al., 1994). Although such hairpin structures were found in a few archaeal transcript terminators, they are dispensable for transcription termination, as indicated by both *in vitro* and *in vivo* assays (Santangelo et al., 2009; Santangelo and Reeve, 2006; Thomm et al., 1994). Whereas, the terminator U-tracts, usually comprised of 4–7 consecutive uridines, are essential for efficient termination, as shown using an *in vitro* transcription system as well as an *in vivo* reporter system in *T. kodakarensis*, in which the tested terminators were inserted just +10 nt downstream from the transcription start sites (Santangelo et al., 2009; Santangelo and Reeve, 2006; Thomm et al., 1994). Nevertheless, the *in vitro* assay may not yet distinguish whether the terminator U-tract merely caused TEC pausing or also dismantled it from the DNA template. While putting terminator closely downstream the transcription start site in the *in vivo* reporter system could introduce artificial assay biases and may not tell the virtual *in vivo* transcription termination action, as TEC has not yet transited into the normal transcription elongation status and abortive transcriptions usually occur at the transcription initiation phase caused by RNAP reiterative initiation (Blombach et al., 2019; Smollett et al., 2017).

This work, through analyzing the Term-seq data, for the first time, systematically quantified the contributions of both the terminator U-tracts and the termination factor aCPSF1 to *in vivo* TTEs at the genome-wide level, and in combination with extensive biochemical, genetic, and molecular assays, our results indicated that neither aCPSF1 nor the terminator U-tract is dispensable and the two synergistically function in effective and hierarchical transcription termination of *M. maripaludis*. The genome-level Term-seq analysis indicated that both the terminator U4-tracts and the aCPSF1 dependency have a linear positive correlation with the TTEs of 961 identified TTSs of TUs (Figures 1 and 2) and 76.6% (736/961) TUs were determined having both U4-tract terminators and aCPSF1 dependency (Figure 1 - figure supplement 4A), and also revealed that terminators containing more U4-tracts numbers have higher aCPSF1 dependency (Figure 2C and 2D). Accordingly, the *in vitro* biochemical assays clearly demonstrated that aCPSF1 binds more strongly to the terminators containing more U-tracts and two U4-tracts could be the minimum specific sequence for efficient binding of aCPSF1 (Figures 3, 4, Figure 3 - figure supplement 1, 2 and Figure 4 - figure supplement 1). Meanwhile, the terminator reporter assays further verified the synergism of aCPSF1 and the terminator U-tracts in dictating high TTEs *in vivo* (Figure 5). Therefore, based on these results, we argue that the termination factor aCPSF1 specifically and hierarchically binds to the terminator U-tract signals embedded in nascent transcript 3′-ends, and synergistically functions with them to dictate the genome-wide TTEs in *M. maripaludis*. Given the wide distributions of both aCPSF1 and the terminator U-tracts in the representative archaeal species (Berkemer et al., 2020; Dar et al., 2016a; Yue et al., 2020), we propose that the synergistic action mode of aCPSF1 and terminator U-tracts could be widely employed by archaeal phyla and may represent the primary archaeal transcription termination mode. Additionally, aCPSF1 was also found guaranteeing the termination of 15% (144/961) TUs with weak terminator signals lacking U4-tract in *M. maripaludis* (Figure 1 - figure supplement 4A), which provides the first evidence for the hypothesis that aCPSF1 could play a back-up role for terminating the transcripts with less efficient intrinsic terminators. Therefore, by collaborating with the intrinsic terminator U-tracts for high TTEs and back-up for weak terminators without U-tract, aCPSF1-dependent termination could serve as the principal termination mechanism in Archaea.

### The N-terminal KH domains equip aCPSF1 to bind the terminator U-tract and function in transcription termination *in vivo*

The resolved structures of aCPSF1 orthologs from *M. thermautotrophicus* (Silva et al., 2011), *M. mazei* (Mir-Montazeri et al., 2011) and *Pyrococcus horikoshii* (Nishida et al., 2010) all carry two KH domains (KHa and KHb) at the N-terminus, which are assumed to be the RNA binding module (Phung et al., 2013; Silva et al., 2011). Since its identification (Siomi et al., 1993), the KH domain, either as single or multiple copies, has been widely found in proteins with various cellular functions, and it has been assumed to be a nucleic acid recognition module and essential to the proteins to bind their specific targets. KH domains commonly occur in proteins involving in transcriptional, posttranscriptional and translational regulation, and other cellular processes. E.g. the KH domains of the IMP family of mRNA binding proteins (ZBP1/IMP1, IMP2, and IMP3) play roles in targeting unique pools of RNAs and so equipping the proteins involved in regulation of RNA stability, localization, and translation (Biswas et al., 2019). The two KH domains of NusA are key for the protein to play regulatory roles in transcriptional elongation, pausing, termination and antitermination in bacteria (Beuth et al., 2005; Gibson et al., 1993). The KH/S1 domains of the eukaryotic exosome complex comprise the central channel for passing divergent RNA substrates, and are thus involved in RNA editing, processing, quality control, or turnover (Wasmuth et al., 2014), while the KH/S1 domains of the bacterial polynucleotide phosphorylase (PNPase) play critical roles in RNA substrate binding and autoregulation (Wong et al., 2013).

The presence of KH domains restricted in the aCPSF1 orthologs among the archaeal β-CASP family ribonucleases suggests that aCPSF1 could target more specific RNA sequences than other β-CASP proteins, like aRNase J and aCPSF2, which only contain the MβL and β-CASP domains. Fluorescence polarization anisotropy assays have determined that *M. thermautotrophicus* aCPSF1 (MTH1203) interacts specifically with U-rich oligonucleotides, U7, U4C4, and C4U4 at micromolar affinities, but not with C7, A7 or G7 oligonucleotides, while the specific interaction was not detected between its N-terminal KH domain and the U-rich oligonucleotides (Silva et al., 2011). However, in the present study, we determined that the KH domains of *M. maripaludis* aCPSF1 exhibit a binding ability and specificity to native terminator U-rich sequences preceding TTSs similar to the whole-length protein (Figures 3, 4 and 6A), therefore demonstrating that the N-terminal KH domains of aCPSF1 are the functional module for specifically recognizing the U-rich terminator signals embedded in transcript 3′-ends. Complementation with aCPSF1 lacking the KH domains in ▽*aCPSF1* cannot restore the growth and transcription termination defects (Figures 6B–D and Figure 6 - figure supplement 1), further supporting that deletion of KH domains led aCPSF1 to lose the capability of mediating transcription termination as being incapable of recognizing the U-rich terminators. The necessity of U-tract recognition in dictating TTEs was further confirmed by the *in vivo* terminator reporter assays, and the correlation between the terminator U4-tracts numbers and TTEs was much weaker in the KH deletion complementary strain Com(ΔKH) than in the wild-type strain, being at a similar level as in the aCPSF1 depletion strain (▽*aCPSF1*) (Figure 5). Taken together, the *in vitro* and *in vivo* experiments prove that the two N-terminal KH domains equip aCPSF1 with the ability of specifically recognizing and binding the terminator U-tract sequences and such recognition is vital for aCPSF1 to function in transcription termination *in vivo*.

### aCPSF1-dependent archaeal transcription termination could represent a simplified eukaryotic RNAP II termination model by a single protein implementing specific binding to terminator signals and cleavage of transcript 3′-ends

Based on the findings in this work, we propose that the multi-functional domain aCPSF1 may perform both terminator signal recognition and transcript 3′-end cleavage to trigger archaeal transcription termination by itself (Figure 7). First, the intrinsic U-tract terminator signals at nascent transcript 3′-ends could be recognized by the N-terminal KH domains of aCPSF1, and direct association of aCPSF1 with aRNAP provides the spatial convenience for the terminator signal recognition. Initiated by such specific binding, the nuclease module of aCPSF1 could perform the downstream endoribonucleolytic cleavage to produce the transcript 3′-end, which has also been determined to be necessary for transcription termination (Yue et al., 2020) and the U-tract mediated hierarchical TTEs (Figure 5B) *in vivo*. The U-tract terminator could also cause aRNAP pausing and disturb the TEC, so further synergistically contributing to transcription termination. Therefore, the termination factor aCPSF1, through recognizing and specifically binding to the terminator U-tracts and cleaving at the transcript 3′-ends, in synergy with U-tract intrinsic terminator signal dictates the genome-wide TTEs in Archaea.

The archaeal aCPSF1-dependent termination mechanism involving transcript 3′-end cleavage has exposed the resemblance with the eukaryotic 3′-end processing/cleavage triggered RNAP II termination mode, in which an aCPSF1 homolog, the yeast Ysh1 or human CPSF73, in the multi-subunits composed 3′-end processing machinery executes the 3′-end cleavage (Dominski, 2010; Eaton et al., 2018; Hill et al., 2019; Kuehner et al., 2011; Porrua et al., 2016). However, despite being homologs, the archaeal aCPSF1 protein carries the N-terminal KH domains to perform the terminator signal recognition by itself, while the eukaryotic CPSF73/Ysh1 have no KH domains and lack the capability of termination signal recognition, which is dependent on the accessory factors. The interacting subunits in the CPF/CPSF complex and the accessory cleavage factors IA and IB (CFIA and CFIB) recognize the poly(A) terminator signal and the flanking U-rich sequences to facilitate the 3′-end cleavage by Ysh1/CPSF73 (Hill et al., 2019; Porrua and Libri, 2015). Thus, the transcript 3′-end cleavage mode and the U-rich sequence recognition by the RNAP II termination machinery could be the evolutionary relic retained in the eukaryotic termination mechanism. However, compared to the simplified archaeal termination mode, which is mainly dependent on one single factor (aCPSF1), the eukaryotic termination process is complex and multi-faceted often involving multiple trans-acting (termination) factors and intertwined with RNA processing/maturation event. Very few homologs of the eukaryotic 3′-end processing complex (Casanal et al., 2017; Shi et al., 2009) and termination factors (Baejen et al., 2017; Eaton et al., 2018; Grzechnik et al., 2015; Larochelle et al., 2018) have been identified in Archaea, and no presumed accessory homologs were found in the aCPSF1 co-immunoprecipitated proteins in our preliminary study. Noteworthily, the eukaryotic CPSF73-mediated cleavage downstream from the 3′-end poly(A) site not only terminates transcription but also provides a polyadenylation site for transcript maturation (Dominski, 2010; Yang et al., 2009), whereas no polyadenylation follows aCPSF1 3′-end cleavage in Archaea. Therefore, we propose that the aCPSF1-dependent archaeal termination mechanism reported here could be a simplified and evolutionary predecessor of the eukaryotic transcription termination machinery.

In conclusion, this work reports that the intrinsic terminator U-tract cis-element and the trans-acting termination factor aCPSF1 synergistically dictate the effective and hierarchical transcription termination efficiency in Archaea at the genome-wide level. The two-in-one mechanism, but not two independent termination systems, could be the principal termination mechanism of Archaea. Although resembling the eukaryotic RNAP II termination mode, Archaea could employ a simplified termination strategy, as aCPSF1, a single factor, could initiate archaeal transcription termination through (i) specifically binding of the N-terminal KH domains to the terminator U-tract and (ii) cleavage at the transcripts 3′-end by the nuclease domain. Therefore, the archaeal transcription termination mode might represent the archetype of the eukaryotic RNAP II transcription termination.

## Materials and Methods

### Strains, plasmids and culture conditions

Strains and plasmids used in this study are listed in Table S1. *M. maripaludis* S2 and its derivatives were grown in pre-reduced McF medium under a gas phase of N_2_/CO_2_ (80:20) at 37°C and 22°C as previously described (Sarmiento et al., 2011), and 1.5% agar was used in the solid medium. Neomycin (1.0 mg/ml) and puromycin (2.5 µg/ml) were used for genetic modification selections unless indicated otherwise. *E. coli* DH5α, BL21(DE3)pLysS and BW25113 were grown at 37°C in Luria-Bertani (LB) broth and supplemented with ampicillin (100 μg/ml), streptomycin (50 μg/ml) or kanamycin (50 μg/ml) when needed.

### Term-seq data analysis

TTS analysis was based on the Term-seq data obtained in previous study based on two parallel replicates (Yue et al., 2020). To minimize the possibility of sites derived from stale RNA processing or degradation products, TTSs were all assigned within 200 nt downstream the stop codon of a gene to maximumly enrich the authentic TTSs near the gene 3′-ends and only sites that appeared in the two replicates with significant coverage were recorded. The primary TTSs were identified as described previously (Yue et al., 2020). Other TTSs except the primary TTSs were assigned as the secondary TTSs (Datasheet S1) by fitting the following three criteria: (i) within 200 nt downstream the stop codon of a gene; (ii) >1.1 read ratio of −1 site (predicted TTS) to +1 site (1 nt downstream TTS); (iii) read-counts of −1 site minus +1 site >5. Motif sequence logos were created using WebLogo (http://weblogo.threeplusone.com/) developed by Schneider and Stephens (Crooks et al., 2004). The lengths of 3′UTR were defined as the distance between the stop codons to the TTSs and listed in Datasheet S1. Number of U4 tracts in the range from −31 site (30 nt upstream the TTS) to −1 site (identified TTS) of each primary TTSs was searched and listed in Datasheet S2.

Transcription termination efficiency (TTE) for transcript with Term-seq identified TTS was calculated by the formula as follow,

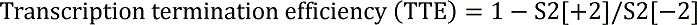

where S2[+2] and S2[-2] indicate the read-counts of +2 (2 nt downstream TTS) and −2 site (1 nt upstream TTS) in strain S2, respectively. Next, TTS Quadruplet Read Ratio (TQRR) was developed to define the transcription termination dependency on aCPSF1 using the following equation:

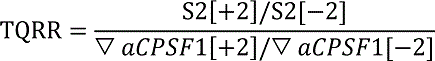

where S2[+2] and S2[-2] indicate the read-counts of +2 and −2 site in strain S2, respectively; ▽aCPSF1[+2] and ▽aCPSF1[-2] indicate the read-counts of +2 and −2 site in strain ▽aCPSF1, respectively. Based on the criteria, those with TQRR < 1 were defined as aCPSF1-dependent terminators.

### Rapid amplification of cDNA 3′-ends (3′RACE)

3′RACE was performed as described previously (Zhang et al., 2009). Total RNA was extracted from the cells cultured to the exponential phase, and a total of 20 μg were ligated with 50 pmol 3′adaptor Linker (5′-rAppCTGTAGGCACCATCAAT–NH_2_-3′; NEB) via a 16 h incubation at 16°C using 20 U T4 RNA ligase (Ambion). Isopropanol precipitation was performed for the 3′ linker-ligated RNA recovery; one aliquot of 2 μg recovered 3′ linker-ligated RNA was mixed with 100 pmol 3′R-RT-P (5′-ATTGATGGTGCCTACAG-3′, complementary to the 3′ RNA linker) by incubation at 65°C for 10 min and on ice for 2 min. Then, the reverse transcription (RT) reaction was performed using 200 U SuperScript III reverse transcriptase (Invitrogen). After RT, nested PCR was conducted using primers (Table S2) that target regions of <200 nt upstream the termination codons to obtain gene-specific products. Specific PCR products were excised from 2% agarose gel and cloned into pMD19-T (TaKaRa) and then sequenced. The 3′-end of a TU is defined as the nucleotide linked to the 3′RACE-linker.

### Protein purification of aCPSF1 and its N-terminal KH domain

The heterogeneous expression and purification of *Mmp-*aCPSF1 in *E. coli* have been described in detail previously (Yue et al., 2020). For purification of the aCPSF1 KH domains, the 1-147 amino acids of *Mmp*-aCPSF1 including the N-terminal two KH domains was inserted into pET28a using the ClonExpress MultiS One Step Cloning Kit. The constructed plasmid was then transformed into *E. coli* BL21(DE3)pLysS to heterogeneously express a His-tagged aCPSF1-KH domain recombinant protein. After a 16-h induction at 22°C, the cells were harvested, and the protein was purified through a His-Trap HP column and followingly a Q-Trap HP column as described previously (Zheng et al., 2017). Purified proteins were detected by SDS-PAGE, and the protein concentration was determined using a BCA protein assay kit (Thermo Scientific).

### RNA Electrophoretic Mobility Shift Assay (rEMSA)

The rEMSA assay was performed as described previously with some modifications (Li et al., 2019). In Brief, an RNA binding assay was performed in a 10 μl reaction mixture containing binding buffer (20 mM HEPES-KOH, pH 7.5, 1 mM MgCl_2_, 150 mM NaCl, and 5% of glycerol), 0.5-nM 3′-biotin-labeled RNA substrate, and increasing amounts of purified recombinant proteins of the catalytic inactive aCPSF1 and its N-terminal KH domains (aCPSF1-KH). After incubation at 25°C for 30 min, the reaction mixtures were loaded onto a 6% polyacrylamide gel and electrophoresed under 100 V for 50 min in 0.5 × TBE running buffer. The free RNA and RNA-protein complexes in gels were transferred onto a nylon membrane and cross-linked using GS Gene Linker™ UV Chamber (Bio-Rad Laboratories, Hercules, CA). The nylon membrane was incubated with 20 μg/mL of proteinase K (Ambion) at 55°C for 2 h, then a Chemiluminescent Nucleic Acid Detection Module kit (Thermo Scientific) was used to detect chemiluminescence by exposure on a Tanon-5200 Multi instrument (Tanon Science & Technology Co. Ltd., Shanghai, China).

### Surface Plasmon Resonance (SPR) Assay

A surface plasmon resonance (SPR) assay was performed to determine the binding affinities of aCPSF1 to poly-U-tract containing RNA on a BIAcore 8K instrument (GE Healthcare) as described previously (Zhang et al., 2017) with minor modifications. A streptavidin-coated sensor chip SA (Series S Sensor chip SA®, GE Healthcare) was first conditioned with three injections (10 µl min^−1^) of buffer containing 1 M of NaCl and 50 mM of NaOH until a stable baseline was obtained.

The 3′-biotinylated RNA was then diluted to 200 nM in binding buffer (20 mM HEPES-KOH, pH 7.5, 1 mM MgCl_2_, 150 mM NaCl, 5% of glycerol, and 0.05% Tween 20) and immobilized in flow cell 2 at a flow rate of 10 µl min^−1^ for 5 min.

NaCl (500 mM) was then injected at 5 μl min^−1^ to remove unbound RNA molecules until the response units (RU) reached a stable state. *Mmp*-aCPSF1 was twofold serially diluted from 1000 to 0 nM with binding buffer and continuously injected into flow cells with RNA immobilized and the control flow cell 1 without RNA of the sensor chip simultaneously at room temperature. The signal of flow cells with RNA was subtracted from that of flow cell 1 to eliminate nonspecific interactions. BSA was included as a negative control. The sensorgrams were analyzed using Biacore Insight Evaluation Software (version 1.0.5.11069, GE Healthcare).

### Construction of the ΔKH-aCPSF1 complementary strain

The *Mmp-aCPSF1* (*MMP0694*) expression depleted strain (▽*aCPSF1*) was constructed using a *TetR*-*tetO* repressor-operator system as described previously (Yue et al., 2020). To obtain an aCPSF1-ΔKH complementary strain, the plasmid pMEV2-*aCPSF1*-ΔKH was first constructed. To obtain this plasmid, the fragment *aCPSF1*ΔKH-pMEV2 was amplified from the plasmid pMEV2-*aCPSF1* using primers pMEV2-*aCPSF1*-ΔKH-F/R. Through the Gibson assembly via ClonExpress Ultra One Step Cloning Kit (Vazyme), the fragment *aCPSF1*ΔKH-pMEV2 was then circled to form the plasmid pMEV2-*aCPSF1*-ΔKH. The constructed plasmid was transformed into ▽*aCPSF1* via the PEG-mediated transformation approach to produce the complementary strains (Tumbula et al., 1994).

### Western blot assay

Western blot was performed to determine the cellular *Mmp*-aCPSF1 or aCPSF1-ΔKH protein abundances in various genetic modified strains as described previously (Yue et al., 2020). A polyclonal rabbit antiserum against the purified *Mmp*-aCPSF1 protein was raised by MBL International Corporation, respectively. The mid-exponential cells of *M. maripaludis* were harvested and resuspended in a lysis buffer [50 mM Tris-HCl (pH 7.5), 150 mM NaCl, 10 (w/v) glycerol, 0.05% (v/v) NP-40], and lysed by sonication. The cell lysate was centrifuged and proteins in the supernatant were separated on 12% SDS-PAGE and then transferred to a nitrocellulose membrane. The antisera of anti-*Mmp*-aCPSF1 (1: 10,000) were diluted and used respectively, and a horseradish peroxidase (HRP)-linked secondary conjugate at 1:5,000 dilutions was used for immunoreaction with the anti-*Mmp*-aCPSF1 antiserum. Immune-active bands were visualized by an Amersham ECL Prime Western blot detection reagent (GE Healthcare).

### Construction of the *in vivo* terminator reporter system

To test the TTE of each terminator *in vivo*, the terminator reporter system was constructed by putting the tested terminator between the upstream luciferase and downstream mCherry gene. The promoter of *MMP1697* was used to transcribe this transcript because this promoter has similar transcription ability in strains S2 and ▽*aCPSF1*. To obtain this reporter system, we first synthesis the DNA fragment P1697-luciferase-tested terminator**-**mCherry-T0952, in which P1697 and T0952 are the promoter of *MMP1697* and the terminator of *MMP0952*; luciferase and mCherry are the two codon-optimized fluorescent reporter genes. For the homologous recombination of the synthesized fragment to the *M. maripaludis* genome, the genome region of the intergenic region (IGR) between *MMP0852* and *MMP0853* was chosen as this IGR is long enough to avoid polar effect to downstream genes. Then, to get the homologous arms, the DNA fragment from the upstream of *MMP0852* to the downstream of *MMP0853* was PCR amplified from the S2 genomic DNA using primers *MMP0852up-F* and *MMP0853down-R*, and inserted to pMD19-T (TAKARA) to obtain pMD19-T-MMP0852/0853 (Table S1). The synthesized DNA fragment of P1697-luciferase-Reporter terminator**-**mCherry-T0952 and the resistance gene were inserted into the IGR between *MMP0852* and *MMP0853* via ClonExpress Ultra One Step Cloning Kit to get the plasmid pReporter-Ter. Finally, this reporter system was transformed into *M. maripaludis* S2 and its derivatives via the PEG-mediated transformation approach to produce the reporter system strains (Tumbula et al., 1994). Different terminator sequence fragments were changed through Gibson assembly technique by the primer listed in Table S2.

### Quantitative RT-PCR for transcription abundance determination of reporter genes

Total RNA was extracted from the mid-exponential cells as described previously (Yue et al., 2020). 500 ng Total RNA were digested and reverse transcribed using ReverTra Ace® qPCR RT Master Mix with gDNA Remover (Toyobo) according to the supplier′s instructions and used for qPCR amplification with the corresponding primers (Table S2). Amplifications were performed with a Mastercycler ep realplex2 (Eppendorf AG, Hamburg, Germany). To estimate copy numbers of the luciferase and mCherry mRNA, a standard curve of their genes was generated by quantitative PCR using 10-fold serially diluted PCR product as the template. The number of copies of mCherry transcript per luciferase transcript copies is shown. All measurements were performed on triplicate samples and repeated at least three times.

## Acknowledgments

The authors thank Prof. William B. Whitman at the University of Georgia providing strain *Methanococcus maripaludis* S2 and plasmids pIJA03 and pMEV2. The authors also thank Zheng Fan for surface plasmon resonance assay and Jingfang Liu for recombinant protein identification by MS, and LetPub (www.letpub.com) for linguistic assistance on the manuscript preparation.

## Author contributions

J. L. and L.Y. designed and performed the genetic and biochemical experiments. J.L. and X.Z.D. conceptualized the experiments and acquired funding. L.Y., W.T.Z., Z.H.L., B.Z. and F.Q.Z. performed the Term-seq data analysis. W.T.Z. constructed the terminator reporter transformed strains and Z.H.L. performed part of rEMSA assays. J.L. and X.Z.D. wrote the manuscript and all of the authors approved the final manuscript.

